# The acute effects of cocoa flavanols on cognitive control and response inhibition: A randomised crossover trial

**DOI:** 10.1101/2024.03.10.584219

**Authors:** Ahmet Altınok, Monica Moruzzi, Elkan G. Akyürek

**Author notes:** Address correspondence to: Ahmet Altınok, Department of Experimental Psychology, University of Groningen Grote Kruisstraat 2/1, 9712 TS Groningen, The Netherlands.

## Abstract

In this pre-registered study, we investigated the effects of acute cocoa flavanol (CF) consumption on cognitive control and response inhibition processes, at two different dosage levels. This study was randomised, placebo-controlled, gender-balanced, double-blind, and utilised a crossover design. Participants consumed three different drinks across three separate sessions: A placebo drink with alkalised cocoa powder, a low dosage (415 mg), and a medium dosage (623 mg) of cocoa flavanols from flavanol-rich cocoa powder. Following the administration of these treatment conditions, participants were tested in the Flanker, Simon, and Go/No-go tasks in a counterbalanced order in each session. We analysed accuracy and response times from incongruent and congruent trials of the Simon and Flanker tasks, and commission errors, omission errors, and response times for the Go/No-go task. In addition to these main measurements, we considered interference and sequence effects, accounting for the influence of previous trials in Simon and Flanker tasks. The acute effects of CF on cognitive control and response inhibition were examined using (Generalised) Linear Mixed Model analysis, which included random intercepts, fixed effects, and random slopes. Analysis results revealed that neither dose of cocoa flavanols consumption acutely improved accuracies, interferences, errors, or response times in these three tasks. Furthermore, neither the gender of participants nor BMI scores predicted their cognitive control and response inhibition functions in addition to the treatment conditions. Our findings suggest that acute consumption of cocoa flavanols does not significantly enhance cognitive control or response inhibition in healthy young adults.

## Introduction

Cocoa, commonly enjoyed in various forms for its distinct flavour, also contains flavanoids, components found in other foods like red wine, berries, green tea, and grapes, and integral to polyphenols. Flavanols, a specific subgroup of flavonoids within polyphenols, are thought to offer potential health benefits (Andújar et al., 2012; Fraga et al., 2010; Manach et al., 2004; Tsao, 2010). These include reducing the risk of cardiovascular diseases and enhancing overall health and quality of life (Martin & Ramos, 2021).

Cocoa flavanols (CF) have also been shown to enhance cognition by increasing nitric oxide (NO) synthesis, a critical factor in improving cerebral blood flow and neurotransmission. Studies demonstrated CF’s role in elevating NO bioactivity (Heiss et al., 2003; Karim et al., 2000; Loke et al., 2008), which facilitates vasodilation in cerebral arteries (Calver et al., 1992), and strengthens neural communication (Hardingham et al., 2013; Vincent, 2010). This action of NO, particularly in synaptic areas, is crucial for cognitive functions like memory and learning (Epstein et al., 1993; Huang, 1997), underscoring CF’s potential to boost cognitive control. Beyond its bioavailability and psychological benefits, there is an increasing trend of recent randomised clinical on the acute effects of CF in varied doses on various cognitive functions across different age groups. These functions include cognitive control, working memory, and various subtypes of attention, such as spatial, temporal, and sustained attention. However, it is important to note that the results from these trials have presented a varied picture to date.

Regarding working memory and sustained attention, an initial behavioural study reported significant improvement in working memory and sustained attention by administering 530mg and 994mg CF in young adults (Scholey et al., 2010). In this study, Serial subtraction tasks (Serial Threes and Serial Sevens) were used for working memory assessment, which require participants to count backwards from a given number as quickly and accurately as possible. The Rapid Visual Information Processing task (RVIP) task was utilized to evaluate sustained attention, in which the participants were monitoring a continuous sequence of numbers, identifying instances where three consecutive odd or even digits appeared. The 520mg CF dose acutely improved the number of correct responses in the Serial Three task and was overall even slightly better than the 994mg dose; reaction times for the RVIP task were significantly faster in the 520mg CF condition compared to both the 994mg CF and control conditions. Following this, Massee and colleagues (2015) documented positive acute effects of 250mg CF, specifically on the number of correct responses in the Serial Seven task, in slight contrast to Scholey et al. (2010), who observed significant improvements in the Serial Three task. Moreover, Boolani et al. (2017) found that administering 499mg of CF did not lead to improvements in performance on either the Serial Seven or Serial Three tasks.

This inconsistency in CF’s effects extends to other research examining its impact on working memory. Field and colleagues (2011) found significant acute effects of 720mg CF on accuracy in spatial working memory, which involved detecting changes in the memorized locations of objects. Also, in a study by Grassi and colleagues (2016), 520mg CF acutely improved working memory accuracy, measured with the N-back (2-back) task, specifically in the female subgroup in a sleep deprivation condition, while no significant effect was found on sustained vigilance. However, recently, Altınok et al. (2022), in two crossover experiments, reported that 415mg CF did not improve visual working memory maintenance, nor updating. Pase et al. (2013) also reported an absence of effects of 250mg and 500mg polyphenols, the family to which cocoa flavanols belong, on spatial working memory and sustained attention.

Similar inconsistencies remain for the effects of CF on sustained attention. For instance, on RVIP performance, in contrast to Scholey et al. (2010), Massee and colleagues (2015) could not observe any improvement in reaction times or accuracy after consumption of 250mg CF. Boolani et al. (2017) used a dual RVIP, in which a secondary target was added to the task, asking participants to respond to specific numbers (e.g., “5”), in addition to the primary RVIP task. A dose of 499mg CF improved only secondary target response time and overall false alarms. However, there was no improvement in accuracy or reaction times in the primary RVIP task. These findings suggest that while cocoa flavanols may influence certain elements of RVIP performance, their overall effectiveness in enhancing the primary aspects of sustained attention remains inconsistent and not conclusively established. Similar variety was observed in other attention components. Karabay and colleagues (2018) found effects of 374mg and 747mg CF on spatial attention, such that CF significantly decreased reaction time in visual search, but found no improvement in temporal attention.

In terms of cognitive control, Francis and colleagues (2006) tested both acute (450mg) and sub-chronic (172mg for five days) effects of CF on the blood oxygen level-dependent (BOLD) response during task switching in young people. Participants engaged in a task-switching task, alternating between identifying letters (as vowels or consonants) and numbers (as odd or even) among letter-number pairs, based on colour cues during “switch” and “nonswitch” blocks. The authors reported a significant increase in cerebral blood flow and increased activity in the medial and lateral prefrontal cortex, and parietal cortex. These areas were associated with processing response conflict and attention (Rushworth et al., 2001). This improvement in cerebrovascular reactivity aligns with earlier findings on peripheral endothelial function (Heiss et al., 2015). However, despite these neural changes, at the behavioural level, cocoa flavanols did not significantly affect reaction time in switched or non-switched conditions, switching costs, or error rates. Likewise, a functional near-infrared spectroscopy (fNIRS) study (Decroix et al., 2016) investigated the effects of an acute administration of 903mg CF and reported an increase in cerebral oxygenation during the classic Stroop task among twelve young adult males, but, similar to Massee et al. (2015), did not observe significant behavioural changes attributed to CF intake in terms of accuracy or reaction time.

Another fMRI study (Decroix et al., 2019) revealed that 2 hours after intake of 900mg of CF, BOLD signals in the supramarginal gyrus of the parietal lobe and the inferior frontal gyrus were notably enhanced during a Flanker task. This study also demonstrated that such a dosage of CF led to improved cognitive performance, evidenced by a decrease in reaction time compared to the placebo, although this indicated a general treatment effect rather than a specific interaction with the task conditions (e.g., congruent vs. incongruent trials). Another study (Tsukamoto et al., 2018) reported more specific acute effects of 563mg CF on Stroop task interference, on reaction time differences between incongruent and neutral trials. Additionally, the combination of CF and aerobic exercise was also beneficial for reducing interference.

In a recent fNIRS study, Gratton et al. (2020) observed that an acute intake of 682mg of CF enhanced cerebrovascular reactivity in the frontal cortex of healthy young adults during a modified Stroop task. Moreover, the study noted that CF intake specifically boosted cognitive performance in high-demand situations. This was evident during the more complex ‘double Stroop’ task, which introduces dual conflict scenarios – both at the stimulus classification and response selection stages, as opposed to the standard Stroop Task that involves conflict only at the stimulus classification level.

From the studies to date on the acute effects of CF on cognition, the effects found on cognitive control seem the most compelling. Several studies (Decroix et al., 2019; Francis et al., 2006; Gratton et al., 2020; Tsukamoto et al., 2018), have documented significant neural changes, such as increased cerebral blood flow and enhanced cerebrovascular reactivity, particularly in areas of the brain associated with cognitive control functions. Nevertheless, the translation of these neural enhancements into observable behavioural improvements remains inconsistent. Some studies have noted improved cognitive performance following acute CF intake in tasks that demand high cognitive load or involve conflict resolution, like the double Stroop task (Gratton et al., 2020). However, others (Decroix et al., 2016, 2018; Massee et al., 2015; Sasaki et al., 2023, 2024) did not find significant improvements in behavioural outcomes in Stroop task, such as reaction time or accuracy. This disparity suggests that while CF intake may influence brain activity and certain aspects of cognitive control, its effects on measurable behavioural outcomes in cognitive control tasks may depend on the complexity of the task and the specific cognitive demands. The current study was therefore designed to assess different aspects of cognitive control, in particular lower-level cognitive control and response inhibition. In our pre-registered, randomised, placebo-controlled, gender-balanced, double-blind, and crossover-designed trial, we tested the acute effects of two doses (415mg and 623mg) of CF on cognitive control and response inhibition in healthy young adults., using the Flanker Task (Eriksen & Eriksen, 1974), the Simon Task (Simon & Rudell, 1967) and the Go/No-go Task (Drewe, 1975).

Previous studies have predominantly employed the Stroop task and its modifications to assess the effects of CF on cognitive control. The Stroop task is thought to involve interference management and the inhibition of automatic responses (Decroix et al., 2016, 2018; Gratton et al., 2020; Massee et al., 2015; Sasaki et al., 2024; Tsukamoto et al., 2018). The Flanker task has been less frequently used (Decroix et al., 2019), and no studies were found utilizing the Simon task. Despite similarities in addressing interference resolution across the Stroop, Flanker, and Simon tasks, each engages distinct sources of interference and resolution processes. The Stroop task focuses on semantic interference, requiring prioritization of ink colour over word meaning. In contrast, the Flanker Task demands suppression of distracting information to focus on a central target, while the Simon task addresses spatial interference through spatially incongruent stimuli. Yeung and colleagues (2020), using fNIRS, noted weak associations in developmental improvements in inhibition performance between the Stroop and Flanker tasks, underscoring differences in handling spatial and verbal conflicts. Additionally, age-related effects on task performance vary, with older adults showing more significant challenges in the Stroop task than in the Flanker and Simon tasks (Dubravac & Meier, 2023). A study comparing reward effects across these tasks found that while rewards modulate conflict processing, their impact differs, notably not affecting the Stroop effect, but the other two tasks (Mittelstädt et al., 2023). Two ERP studies, highlighted distinct stages of stimuli processing for each task, further differentiated the cognitive control functions reflected by the Flanker N200 and Stroop N450 components (Kałamała et al., 2020; Tillman & Wiens, 2011). It was suggested that the Flanker task might reflect immediate detection of, and response to, spatial conflict, and that the Stroop task might be related to the resolution of conflict that involves more complex cognitive processes, engaged in resolving higher-level, verbal conflict. Mansfield et al. (2013) demonstrated, by means of an ERP analysis, that the Simon and Flanker tasks, though similar, engage different cognitive control mechanisms, with the Simon task involving immediate spatial processing and the Flanker task requiring later, more central processing.

In summary, while the Stroop task is associated with higher-level cognitive processes due to its complex verbal demands, the Simon and Flanker tasks focus on more basic perceptual, peripheral/spatial and motor conflicts. Despite evidence of CF’s impact on higher-level functions, research on its effects on basic cognitive control processes, as measured by the Simon and Flanker tasks, remains scarce. This gap raises questions about the acute influence of CF on lower-level cognitive control functions. In the present study, we therefore implemented both of these tasks.

With regard to response inhibition, Field and colleagues (2011) initially reported reduced reaction times in predictable trials of a sustained attention and response inhibition task, following the administration of 720mg CF. This task required participants to identify presented letters or digits, varying between predictable (only letters, X or Y) and unpredictable (a mix of letters and numbers) conditions. Their findings suggest that acute intake of CF may have the potential to enhance response inhibition in young adults. However, in their research, the task was focused on the sustained attention aspect, more than on response inhibition. In our study, we therefore aimed to directly test the effects of CF on response inhibition more directly, by employing a Go/No-go task. This task is specifically designed to measure inhibitory control, offering a clear measure of the ability to inhibit cognitive processes and control impulsive actions.

## Methods

### Participants

This study involved thirty-six healthy young adults, comprising an equal number of females and males (18 each), within the age range of 19 to 29 years (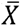= 21.61, SD = 2.61). The study was designed as a crossover randomised controlled trial and based on this, the sample size was calculated using G*Power (Faul et al., 2007), to identify a medium-sized effect. The study parameters included two groups (gender-balanced, females and males) and three measurement points regarding three counterbalanced Treatment conditions. Statistical thresholds were set at α = 0.05, with a power of 0.80, and the critical F-value was calculated as 3.29 (df = 2). The effect size estimation (Cohen’s *d* = 0.325) was adapted from Decroix et al. (2019), who investigated the acute effects of cocoa flavanols on inhibitory control processes in a similar demographic. Participants were presented with an informed consent form, which they were asked to sign before participating in the experiment. This protocol was endorsed by the Ethics Committee of the Psychology Department at the University of Groningen (approval number PSY-2223-S-0149, dated 02.12.2022). The conduct of the study adhered to the ethical standards outlined in the 2008 Declaration of Helsinki.

Participants were required to meet several inclusion criteria: (1) they had no prior diagnosis of any vascular disease and were free from any health conditions affecting metabolism; (2) they were not diagnosed with, or currently suffering from, any neurological or psychiatric disorders and were not following any medically restrictive diets; (3) they did not smoke or use other tobacco products (Francis et al., 2006; Pase et al., 2013), and were free from using over-the-counter or prescription medications, except for contraceptive pills (Scholey et al., 2010), and did not consume vitamin supplements, herbal extracts, or illicit drugs; (4) they were not pregnant or breastfeeding at the time of the study; (5) they possessed normal or corrected-to-normal vision and could perceive colours accurately; (6) their Body Mass Index (BMI) was within the range of 18 to 25.

### General apparatus

Data collection was conducted in the laboratories of the Psychology Department at the University of Groningen. A 27-inch Iiyama Prolite G2773HS monitor was used, featuring a 100 Hz refresh rate and 32-bit colour depth. The monitor’s resolution was set at 1920x 1200 pixels in cognitive task processing. The experimental tasks were designed and executed using OpenSesame 3.3.9 (Mathôt et al., 2012) on a Windows 10 operating system. Responses from participants were recorded via a USB keyboard.

### Experimental products

The study involved two types of cocoa powders: a high-flavanol cocoa powder and an alkalised cocoa powder (see Table 1), both dissolved in 200 ml of tap water and mixed in a 300 ml capacity shaker. Participants consumed three different mixtures, corresponding to three experimental conditions: Placebo, low-level cocoa flavanols, and medium-level cocoa flavanols.

**Table 1.** Nutritional composition of the study Treatments.

| *Experimental ingredients* | **Placebo** | **Low Flavanols** | **Medium Flavanols** |
| --- | --- | --- | --- |
| ***Alkalised cocoa powder (g)*** | **9** | **4** | **1.5** |
| Energy (kcal) | 27.4 | 12.2 | 4.6 |
| Protein (mg) | 1998 | 888 | 333 |
| Fat (mg) | 990 | 440 | 165 |
| Caffeine (mg) | 18 | 8 | 3 |
| Theobromine (mg) | 189 | 84 | 31.5 |
| ***High-flavanol cocoa powder (g)*** | **-** | **5** | **7.5** |
| *Flavanols (mg)* | - | 415 | 623 |
| Energy (kcal) | - | 17.3 | 25.9 |
| Protein (mg) | - | 1120 | 1680 |
| Fat (mg) | - | 700 | 1050 |
| Caffeine (mg) | - | 10 | 15 |
| Theobromine (mg) | - | 105 | 157.5 |
| Water (ml) | 200 | 200 | 200 |

For the placebo (P) condition, 9 grams of alkalised cocoa powder were used. In the low cocoa flavanols (CF-Low) condition, the mixture consisted of 5 grams of high-flavanol cocoa powder (415 mg flavanols) and 4 grams of alkalised cocoa powder. In the medium cocoa flavanols (CF-Medium) condition, 7.5 grams of high-flavanol cocoa powder (623 mg flavanols) and 1.5 grams of alkalised cocoa powder were combined. Alkalised cocoa powder was added in all conditions to standardise the colour and taste of the drinks.

The cocoa powders were supplied by Barry Callebaut company for free, but there was no further involvement in the study. The selected dosages were decided based on the research of Karabay et al. (2018) and Barrera-Reyes et al. (2020), where similar flavanol amounts (374-750 mg) were found to produce significant improvements on cognitive functions.

### General procedure

The experiment was conducted over three separate sessions, utilising a crossover design to ensure all subjects experienced all conditions in counterbalanced sessions: Placebo, low-cocoa flavanols (415 mg), and medium-cocoa flavanols (623 mg). The sequence of Treatments was randomised using a gender-balanced Latin square design. The participants, drawn from a student population at the university, engaged in a variety of academic disciplines. Compensation for participation was offered in the amount of 45 euros or course credits. Comprehensive details about the study, requirements, duration, location, aim, and compensation options, were announced via the university participant pool system to the participants before their participation. Participants were also instructed to abstain from alcohol for 24 hours before each session. To control for the impact of circadian rhythms, testing times were standardised across participants, with the experimental tasks being conducted at 10:00, 11:00, 12:00, and 13:00. Due to the availability of lab cabins, each session could accommodate up to four participants. The intake of experimental beverages occurred 2 h before the cognitive tasks to ensure digestion of flavanols, based on prior research (Francis et al., 2006; Lamport et al., 2015). Following experimental consumption, participants were restricted to water until the commencement of the cognitive tasks. A double-blind methodology was enforced by having different researchers manage drink serving and laboratory instructions, keeping both the lab researcher and the participants blind to the conditions.

In the testing environment, participants were seated in individual cabins, maintaining a consistent screen viewing distance of 60 cm (not fixated). The three computerised tasks administered were aimed at evaluating cognitive control and response inhibition via the Flanker, Simon and Go/No-go Tasks. Task sequences were randomised and counterbalanced across genders. Before starting the tasks, participants were given sufficient time to understand the instructions for the tasks and experimental procedure, and so that the experimenter could clarify any doubts. Additionally, only during the initial session, they completed a brief survey on their habitual intake of caffeine and flavanols, as well as provided demographic data, gender, age, body weight and height. Between sessions, the washout period was 5-9 days.

### Tasks

Figure 1 illustrates three cognitive tasks designed to assess specific aspects of cognitive control and response inhibition. The Flanker and Simon Tasks were employed to evaluate lower-level cognitive control, while the Go/No-go Task was utilised to assess response inhibition.

**Figure 1.**
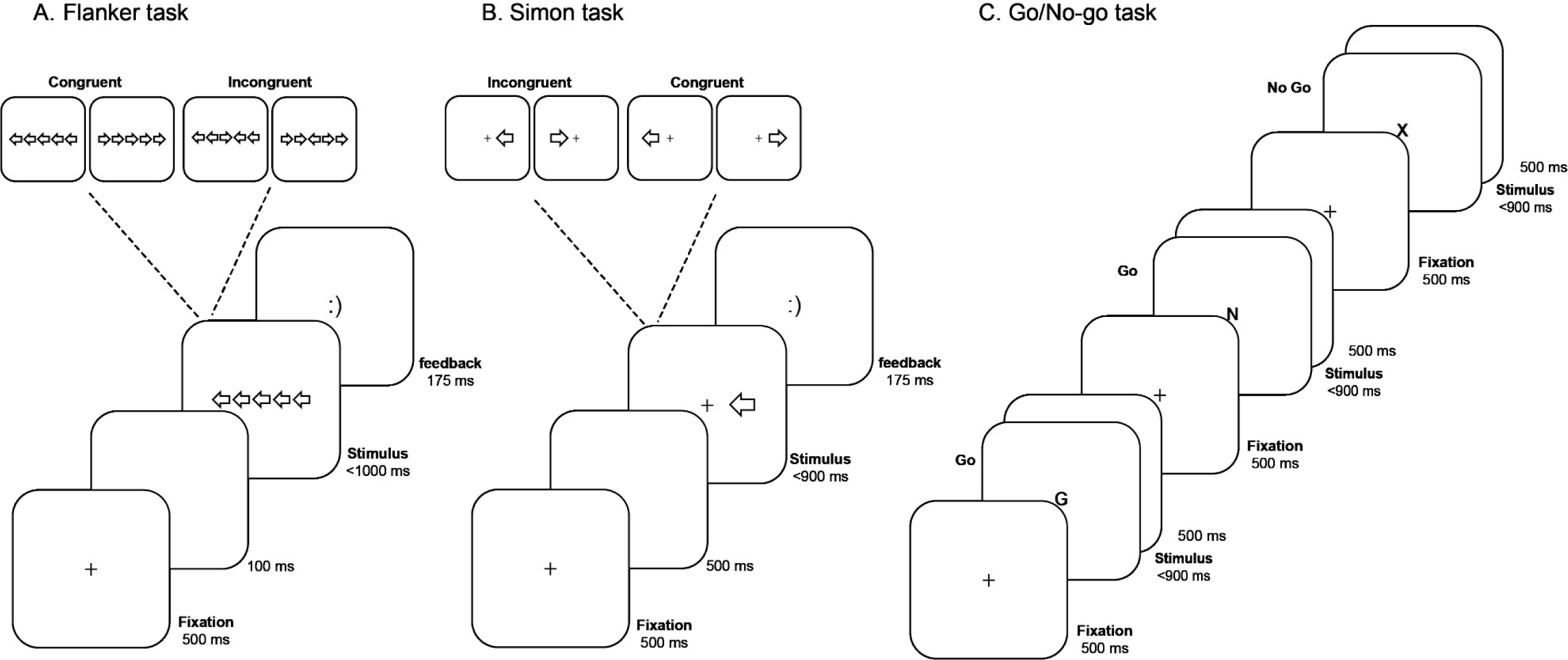
Overview of the Experimental Tasks. *Note*. (a) The Flanker Task is represented, illustrating a congruent trial where the central arrow is surrounded by arrows facing the same direction. (b) The Simon Task is displayed as an incongruent stimulus where the position of the stimulus (right) does not match the arrow’s direction (left). (c) The Go/No-go Task is demonstrated, displaying a sequence that includes a No-go trial, indicated by the “X,” preceded by two Go trials marked by the “G” and “N,” respectively.

The *Flanker task* had four blocks, each containing 100 trials, totalling 400 trials. Before the main experiment, participants were given eight practice trials to familiarise themselves with the task. Within each block, trials were randomised, with 50% of the trials being congruent and the other 50% incongruent. Each trial sequence began with a fixation dot at the centre of the screen for 500 ms, followed by a blank screen for 100 ms, and then the stimulus was displayed. Participants were instructed to identify the direction of the central target arrow, pressing “Z” on their keyboard for left and “M” for right. The arrow display remained visible for 1000 ms or until a response was given. Following the stimuli, feedback was provided for 175 ms, indicating the accuracy of the response. The task stimuli consisted of horizontally oriented arrows, presented in white on a black background. Each arrow had dimensions of 39 pixels in height (equivalent to 2.0° of visual angle, from 60 cm distance) and 60 pixels in width, head to tail, (3.07° of visual angle). The space between arrows was 12 pixels and the total length of the object was 108 pixels, which corresponded to approximately 5.52 degrees of visual angle. The stimuli were arranged in a line of five arrows, with the target arrow located centrally and flanked on either side by two additional arrows. The task comprised both congruent and incongruent trials. In congruent trials, all arrows pointed in the same direction. Incongruent trials featured the target arrow pointing in one direction (left or right) and the flankers in the opposite direction. In this task, the dependent variables (DVs) included accuracy rates, response times for correct responses (RTs), and the Flanker interference effect, which was computed as the difference between RT for incongruent stimuli and RT for congruent stimuli (RT_incongruent_ - RT_congruent_). Additionally, the analysis took into account the trial-to-trial congruency transitions, encompassing sequences such as congruent-congruent (cC), congruent-incongruent (cI), incongruent-congruent (iC), and incongruent-incongruent (iI). In the statistical analysis, responses slower than 100 ms were excluded, leading to the removal of 7 out of 43,200 trials. This exclusion represented 0.02% of the total trials.

Similarly, the *Simon task* had four blocks, each containing 100 trials, totalling 400 trials. Eight practice trials preceded the experimental blocks to familiarise participants with the task. Each block’s trials were randomised, with a balanced distribution of 50% congruent and 50% incongruent trials. Each trial began with the presentation of a central fixation dot, displayed for 500 milliseconds (ms). Following this, the a delay for 500 ms. An arrow pointing to the left or to the right then appeared next to the fixation dot, either 128 pixels from the centre to the left, or to the right side of the screen. The arrow remained on screen for 900 ms, or until a response was registered. If the arrow pointed left, participants were to press the ‘A’ key; if it pointed right, the ‘L’ key would be pressed. After the participants’ responses, feedback was provided for 175 ms, indicating whether the responses were correct or incorrect. The target stimulus, a white arrow displayed on a black background, was 73 pixels high (equivalent to 3.73° of visual angle) and 44 pixels wide (2.25° of visual angle). The task involved congruent trials, where the arrow direction matched its lateral position relative to the fixation dot (leftward arrows pointing left and rightward arrows pointing right), and incongruent trials, where the arrow pointed opposite to its lateral position (leftward arrows pointing right and rightward arrows pointing left). Similar to the Flanker task, the dependent variables (DVs) for this study included accuracy rates for both congruent and incongruent trials, as well as reaction times (RT) for correct responses. In analysing the Simon effect on reaction time, we calculated the difference between the RTs for incongruent and congruent trials (RT_incongruent_ - RT_congruent_). Furthermore, we evaluated the overall probabilities of congruent versus incongruent trials and examined trial-to-trial congruency transitions. These transitions included congruent-congruent (cC), congruent-incongruent (cI), incongruent-congruent (iC), and incongruent-incongruent (iI) sequences. In the statistical analysis, responses registering slower than 100 ms were discarded, excluding 5 out of 43,200 trials, amounting to 0.012% of the total trials.

The *Go/No-go task* commenced with eight practice trials to acclimate participants to the task requirements. Subsequently, a single experimental block containing a randomised sequence of 400 trials was conducted. This block was composed of 270 ‘Go’ trials, accounting for 67.5% of the trials, and 130 ‘No-Go’ trials, constituting 32.5%. Each trial began with a fixation screen lasting for 500 ms, succeeded by the presentation of a letter at the centre of the screen. The letter remained visible for 900 ms, or until a response was given, followed by an interstimulus interval of 500 ms. Participants were presented with ‘Go’ trials, where they were required to respond to the letters ‘W’, ‘R’, ‘Y’, ‘I’, ‘O’, ‘A’, ‘D’, ‘G’, ‘J’, and ‘K’ by pressing the ‘M’ key. Conversely, during ‘No-go’ trials, the letter ‘X’ was shown, and participants were instructed to withhold their response. The target stimuli consisted of 11 letters in Arial font, size 18 pt, each with 24-pixel dimensions corresponding to a visual angle of 1.23 degrees. The stimuli were displayed in black against a grey background (RGB 128, 128, 128) in the centre of the screen. For the Go/No-go task, the dependent variables (DVs) included commission errors (incorrect responses to No-Go trials, difficulty holding not to respond), omission errors (missing responding Go trials), and reaction times (RT) for correct Go trials. During statistical analysis, trials response times below 100 ms were excluded, which was equal to 13 out of 43,200 trials, and represented 0.03% of the total trials.

### Statistical Analysis

The statistical evaluation of the acute impact of cocoa flavanols on cognitive control and response inhibition was carried out using Linear Mixed Models (LMM) and Generalized Linear Mixed Models (GLMM) in RStudio (R Core Team, 2012), with the *nlme* and *lme4* packages (Bates et al., 2015), while the *ggplot2* (Wickham, 2016) and *sjPlot* (Lüdecke, 2022) packages used in the visualisation of results.

Through model evaluation, we compared less and more complex models, focusing on the difference in deviances (Δd) using a chi-square test. The Bayes Factor (*BF_10_*; Wagenmakers, 2007) was calculated based on the Bayesian Information Criterion (BIC). The model tests started with an initial model featuring random intercepts for subjects to evaluate the need for a multilevel model analysis. Subsequently, fixed effects were added to the model, including task conditions (congruent and incongruent for Simon and Flanker tasks), Treatment conditions (placebo, low cocoa flavanols, medium cocoa flavanols), and sessions (1^st^, 2^nd^ and 3^rd^). In analysing the Go/No-go task variables (Commission and Omission errors, response times for correct Go trials), the fixed effects included Treatment conditions and sessions in the model testing.

After the fixed effects analysis, random slopes were included and tested where necessary, and pairwise differences were explored using a Tukey test based on estimated means (LMM) and probabilities (GLMM). In the exploratory analyses, gender and BMI were also considered as fixed effects in the last stage of model testing. Practice trials were excluded from the analysis across all tasks.

In supplement to the multilevel models, Two-Way ANOVA was utilized to probe the Simon effect and Flanker interference on reaction times, including a sequential analysis predicated on previous trial conditions. The analysis tested the main effects of Treatment, Session, and Conditions, which encompassed specific trial sequences—iC (incongruent-congruent), cC (congruent-congruent), iI (incongruent-incongruent), cI (congruent-incongruent)—as well as more general conditions delineated by the congruency of the preceding trial (congruent or incongruent).

### Registration and data availability

The current study was preregistered. The study plan, experimental design, number of participants, cognitive tasks, statistical analysis methods, and data collection procedure are available on the Open Science Framework (https://osf.io/mhnfj). The data of the study and the full analysis scripts are also available on the Open Science Framework (https://osf.io/acvzr).

## Results

### Cognitive Control

#### Flanker Task – Accuracy

In the Flanker Task, the acute effects of CF on accuracy were assessed using a GLMM analysis. Initially, the inclusion of random intercepts for each subject, before adding fixed effects, significantly improved the model 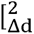 = 372.4, *df* = 1, *p* < .001, *BF_10_* > 100] with deviance changing from 11,752 (BIC = 11763) to 11,380 (BIC = 11401). The addition of fixed effects for the task conditions (congruent and incongruent) led to a significant enhancement of the model 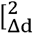 = 408.42, *df* = 1, *p* < .001, *BF_10_* > 100, BIC = 11004]. Still, as fixed effects, the inclusion of session 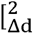 = 11.51, *df* = 2, *p* = .003, *BF_10_* = 0072, BIC = 11013] and Treatment 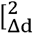 = .39, *df* = 2, *p* = .820, *BF_10_* < .0001, BIC = 11024] did not significantly improve the model. In random slope analysis, a significant improvement was not observed with the addition of random slopes for task conditions 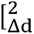 = .13, *df* = 2, *p* = .939, *BF_10_*< .0001, BIC = 11025]. The final model’s fixed and random effects are shown in Table 2, while Figure 2 illustrates the accuracy by condition and Treatment.

**Figure 2.**
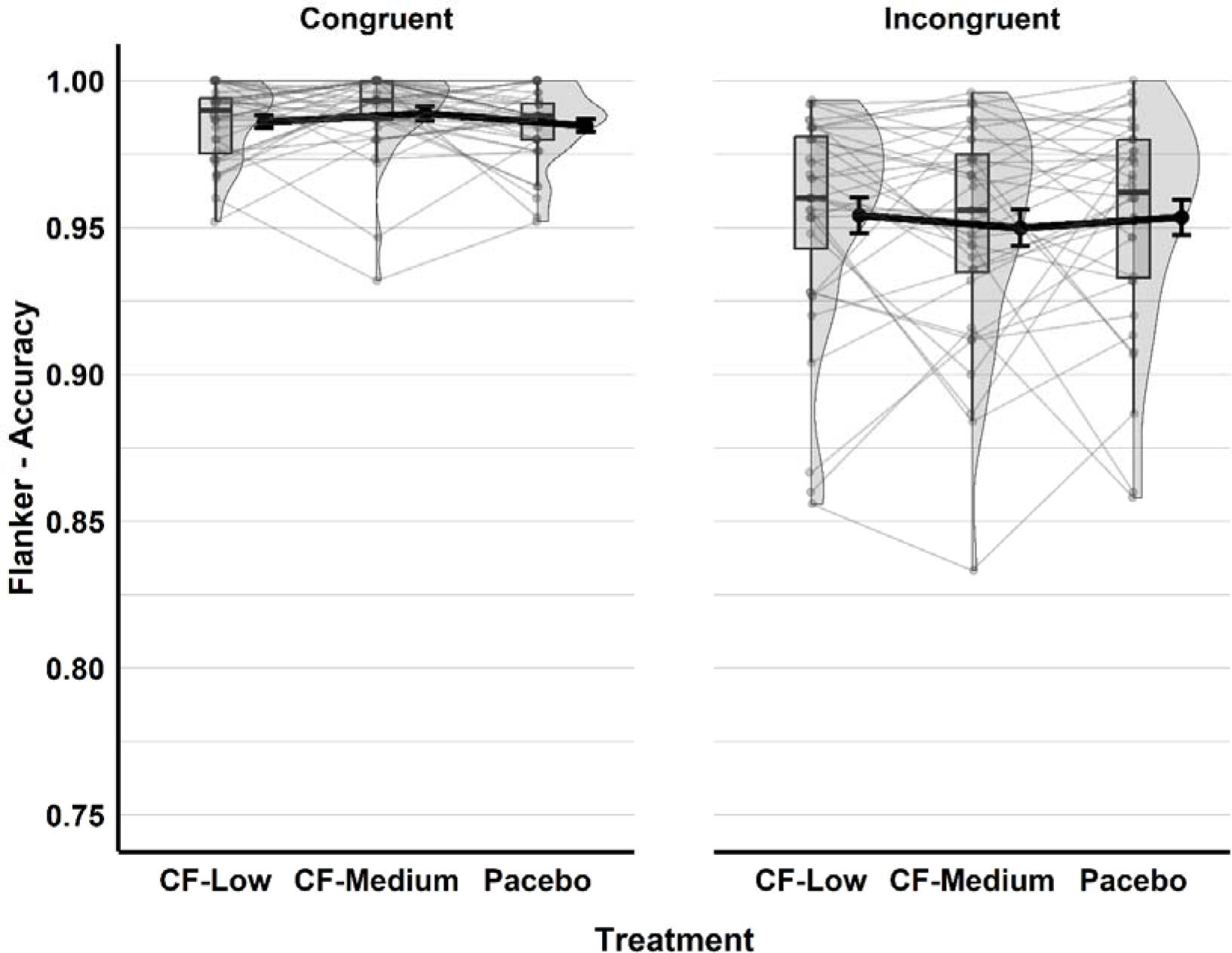
Accuracy in the Flanker Task by Treatment and Condition. *Note*. The boxplots display the data quartiles. Mean values are indicated by black dots with accompanying error bars representing the standard errors. Grey points mark the individual observations, with lines connecting these points between Treatment conditions.

**Table 2.** Generalised Linear Mixed Model Results for the Effects of CF on Flanker Task Accuracy.

| Flanker Task - Accuracy |  |  |  |  |  |
| --- | --- | --- | --- | --- | --- |
| Fixed Effects | *Odds Ratios* | *std. Error* | *CI* | *z* | *p* |
| Intercept | 85.46 | 10.75 | 66.79 – 109.35 | 35.36 | <0.001 |
| Condition [Incongruent] | 0.29 | 0.02 | 0.25 – 0.33 | -18.75 | <0.001 |
| Random Effects | *Variance* | *Sd* |  |  |  |
| τ<sub>00</sub> Subject | 0.44 | 0.66 |  |  |  |
| N<sub>subject</sub> = 36, Observations = 43193 |  |  |  |  |  |

Tukey pairwise comparisons indicated that accuracy in the Flanker task was significantly higher in congruent trials (*prob* = .988, SE = .001, 95% CL [.985 – 0.991]) than in incongruent trials (*prob* = .961, SE = .004, 95% CL [.951 – 0.969]), with Z = 18.75, *p* < .001. In the exploratory analysis, the inclusion of gender 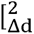 = .09, *df* = 1, *p* = .764, *BF_10_*= .005, BIC = 11014] and BMI scores 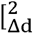 = .77, *df* = 1, *p* = .380, *BF_10_* = .007, BIC = 11013] did not improve the model significantly.

#### Flanker Task – Reaction Time Analysis (General)

The acute effects of CF on response times for correct trials in Flanker task were evaluated with a Linear Mixed Model analysis, and the initial deviance without any fixed or random effects was 491989 (BIC = 492011). Introduction of random intercepts for subjects significantly reduced the deviance to 486933 (BIC = 486965), 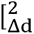 = 5056.5, *df* = 1, *p* < .001, *BF_10_* > 100]. Fixed effects for the task conditions 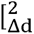 = 1965.7, *df* = 1, *p* < .001, *BF_10_* > 100, BIC = 485010], session 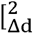 = 952.3, *df* = 2, *p* < .001, *BF_10_* > 100, BIC= 484079] and Treatment 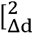 = 25.9, *df* = 2, *p* < .001, *BF_10_* = 46.5, BIC = 484074] resulted in a significant improvement in the model. However, adding the interaction of Treatment and condition 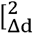 = 2.71, *df* = 2, *p* = .259, *BF_10_* = .002, BIC = 484093] did not improve the model at all.

Following to the fixed effects analysis, the model was significantly improved by adding random slopes for the task conditions 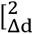 = 83.1, *df* = 2, *p* < .001, *BF_10_*> 100, BIC = 484000] and session 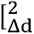 = 706.5, *df* = 7, *p* < .001, *BF_10_* > 100, BIC = 483368] but not Treatment 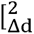 = 13.5, *df* = 11, *p* = .265, *BF_10_* < .0001, BIC = 483471]. Final model results are given in Table 3 and Figure 3 shows the response times for the Flanker task across different conditions and Treatments.

**Figure 3.**
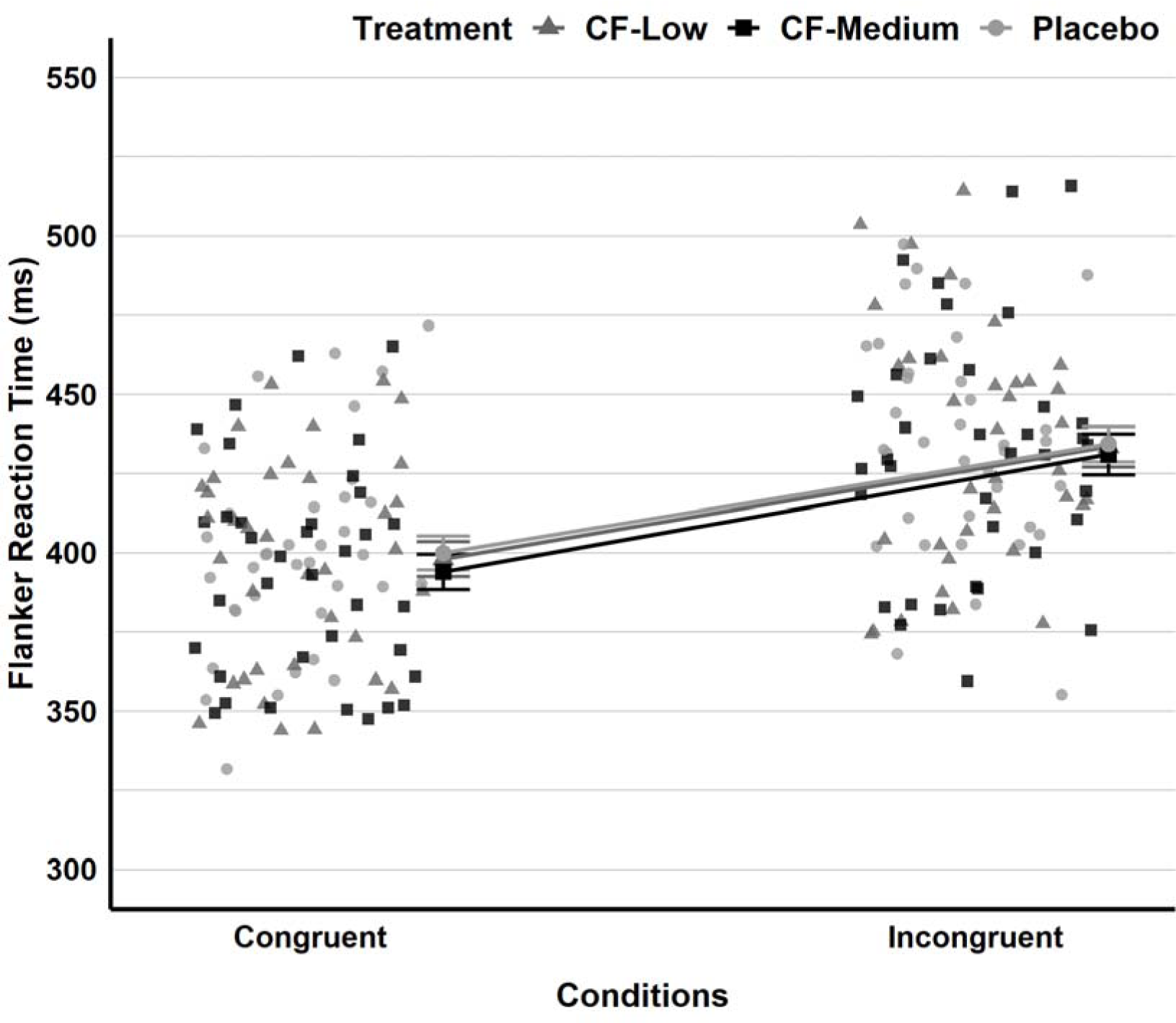
Reaction Time in the Flanker Task Displayed by Treatment and Condition. Note. The Placebo condition is denoted by light grey circles for individual observations and larger dots for means and standard errors. The CF-Low condition is indicated by medium-light grey triangles, while the CF-Medium condition is depicted with black squares.

**Table 3.**
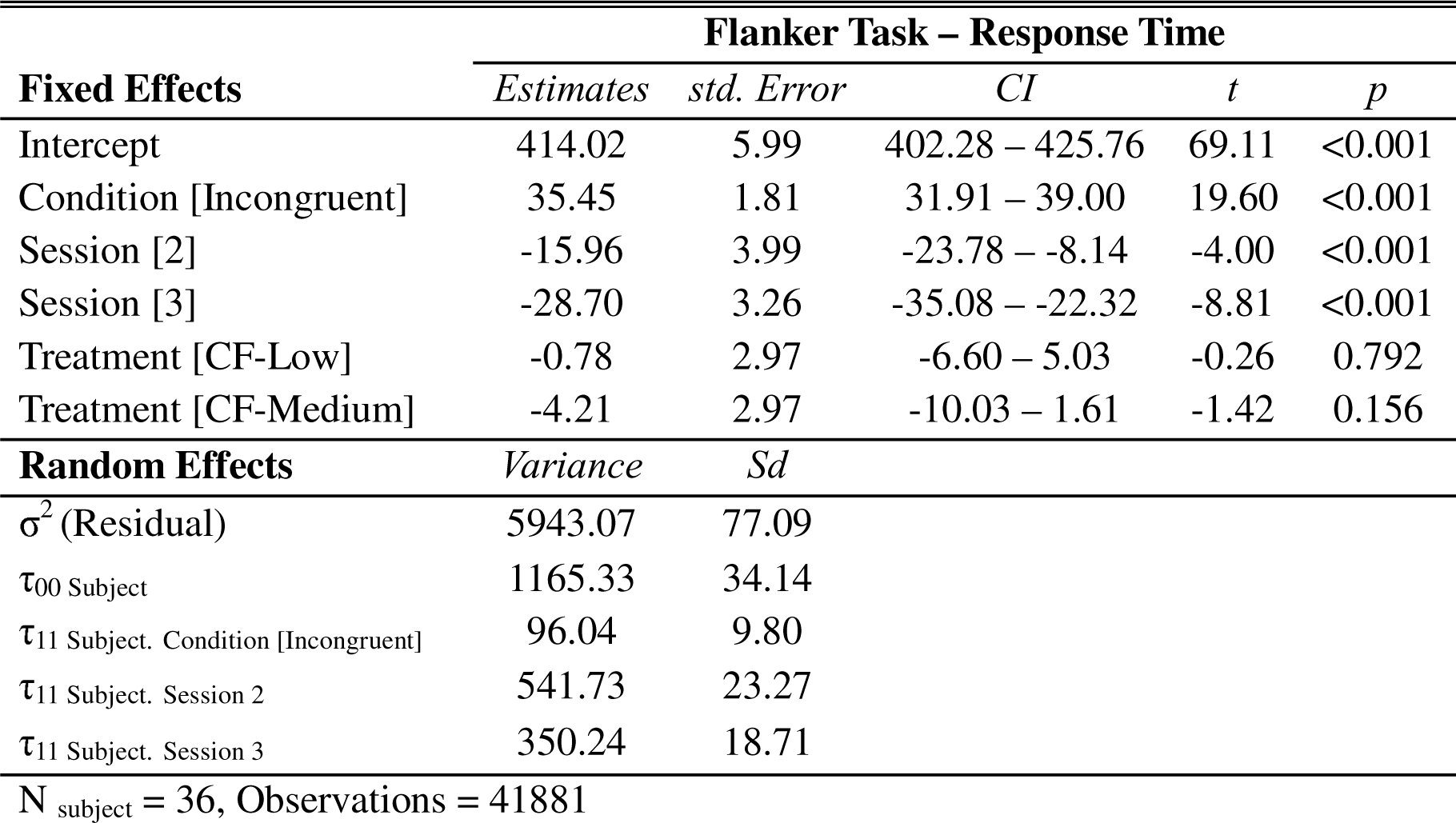
Linear Mixed Model Results for the Effects of CF on Response Time in the Flanker Task.

|  | Flanker Task – Response Time |  |  |  |  |
| --- | --- | --- | --- | --- | --- |
| Fixed Effects | Estimates | std. Error | CI | t | p |
| Intercept | 414.02 | 5.99 | 402.28 – 425.76 | 69.11 | <0.001 |
| Condition [Incongruent] | 35.45 | 1.81 | 31.91 – 39.00 | 19.60 | <0.001 |
| Session [2] | -15.96 | 3.99 | -23.78 – -8.14 | -4.00 | <0.001 |
| Session [3] | -28.70 | 3.26 | -35.08 – -22.32 | -8.81 | <0.001 |
| Treatment [CF-Low] | -0.78 | 2.97 | -6.60 – 5.03 | -0.26 | 0.792 |
| Treatment [CF-Medium] | -4.21 | 2.97 | -10.03 – 1.61 | -1.42 | 0.156 |
| Random Effects | Variance | Sd |  |  |  |
| $\sigma^2$ (Residual) | 5943.07 | 77.09 | | | |
| $\tau_{00}$ Subject | 1165.33 | 34.14 | | | |
| $\tau_{11}$ Subject. Condition [Incongruent] | 96.04 | 9.80 | | | |
| $\tau_{11}$ Subject. Session 2 | 541.73 | 23.27 | | | |
| $\tau_{11}$ Subject. Session 3 | 350.24 | 18.71 | | | |
| N<sub>subject</sub> = 36, Observations = 41881 |  |  |  |  |  |

Pairwise comparisons with the Tukey test indicated that responses were significantly faster for congruent trials (*Estimated marginal mean* = 397, SE = 4.79, 95% CI [388 – 407]) than for incongruent trials (*Estimated marginal mean* = 433, SE = 5.35, 95% CI [422 – 443]), with Z = −19.60, *p* < .001. In the exploratory analysis, the inclusion of gender 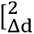 = .78, *df* = 1, *p* = .378, *BF_10_* = .151, BIC = 483401] and BMI scores 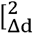 = 1.46, *df* = 1, *p* = .228, *BF_10_* = .034, BIC = 483209] did not significantly improve the model.

#### Flanker Task – Reaction Time Analysis (Sequential)

Besides reaction time analysis within the Flanker task, sequential effects were quantified. The two-way ANOVA results demonstrated significant main effects of previous trial conditions (iC, cC, iI, and cI), *F*(3, 420) = 42.18, *p* < .001, 23. Nonetheless, the main effects of Treatment, *F*(2, 420) = 0.39, *p* = .679, 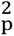 = .002, as well as the interaction between Treatment and conditions, *F*(6, 420) = 0.07, *p* = .999, 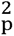 = .001, did not attain statistical significance. Post-hoc analyses employing Tukey’s HSD test revealed that the differences in reaction times between iC and cC trials was not reliable, 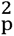 = −4.25, 95% *CI* [−16.80, 8.30], *p* = .819; and likewise, between (cI) and (iI), 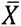 = −8.49, 95% *CI* [−21.03, 4.07], *p* = .303 (see Figure 4).

**Figure 4.**
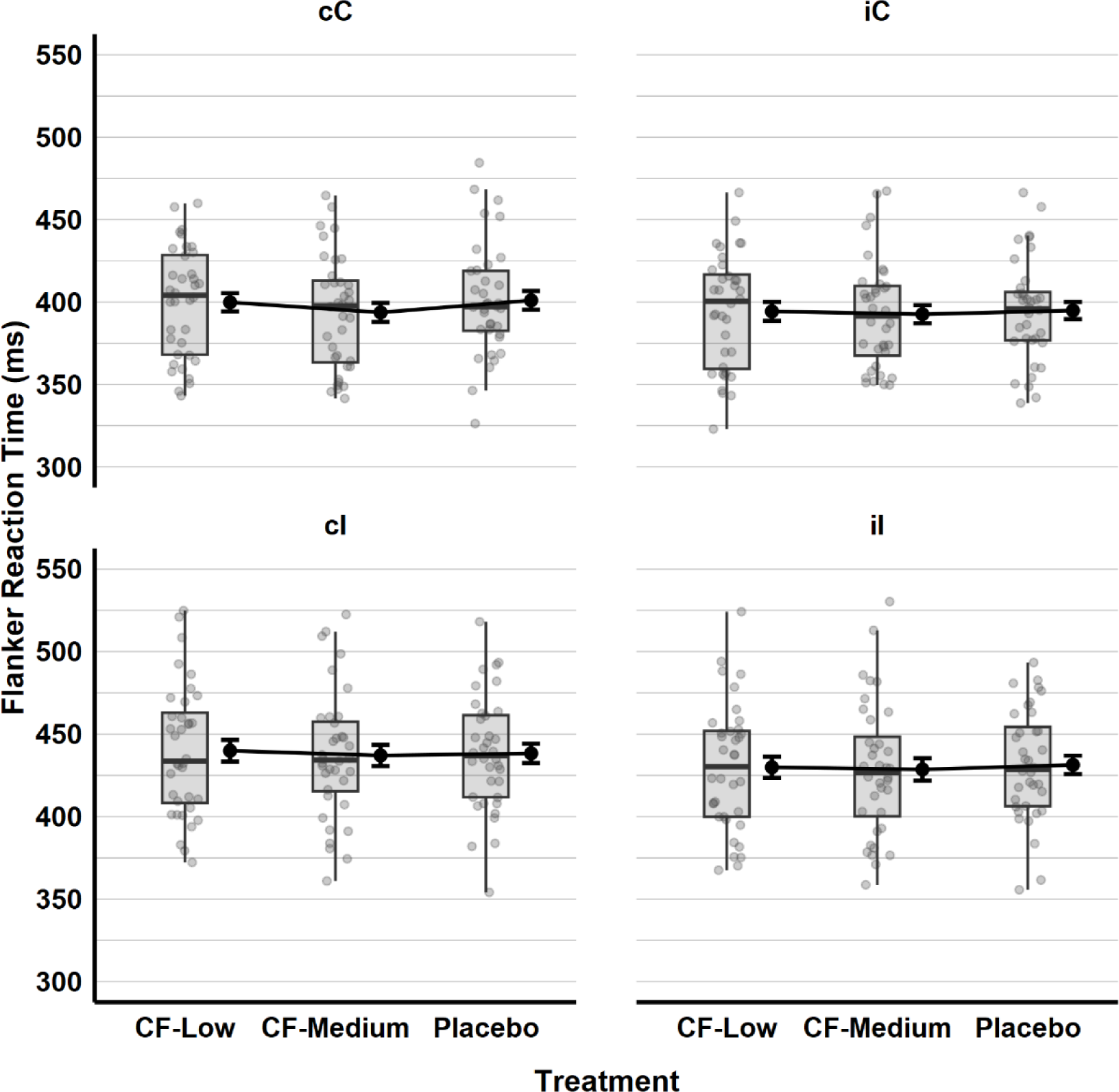
Reaction times in the Flanker Task Across Previous Trial Congruency and Treatment. *Note*. Reaction times in previous (lower case) and current (upper case) trial conditions. The boxplots display the data quartiles. Black dots indicate mean values with accompanying error bars representing the standard errors. Grey points mark the individual observations, in each Treatment conditions.

#### Flanker Interference Effect on Reaction Time (General)

The Flanker effect on reaction time was calculated, and a two-way ANOVA revealed no significant main effects for Treatment, *F*(2, 99) = 0.32, *p* = .73, 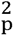 = .006, Session, *F*(2, 99) = 1.02, *p* = .36, 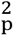 = .02, or the interaction of Treatment and Session, *F*(4, 99) = .56, *p* = .69, 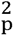 = .02.

#### Flanker Interference Effect on Reaction Time (Sequential)

Similar to the sequential effects of previous trials on general reaction time, the influence of the preceding trial on the Flanker interference effect on reaction time was investigated. A two-way ANOVA indicated that the main effects of previous trial conditions, *F*(1, 210) = 2.70, *p* = .102, 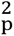 = .01, Treatment, *F*(2, 210) = 0.41, *p* = .667, 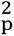 = .004, and the interaction between Treatment and previous trial conditions, *F*(2, 210) = .55, *p* = .580, 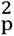 = .005, were not statistically significant (see Figure 5).

**Figure 5.**
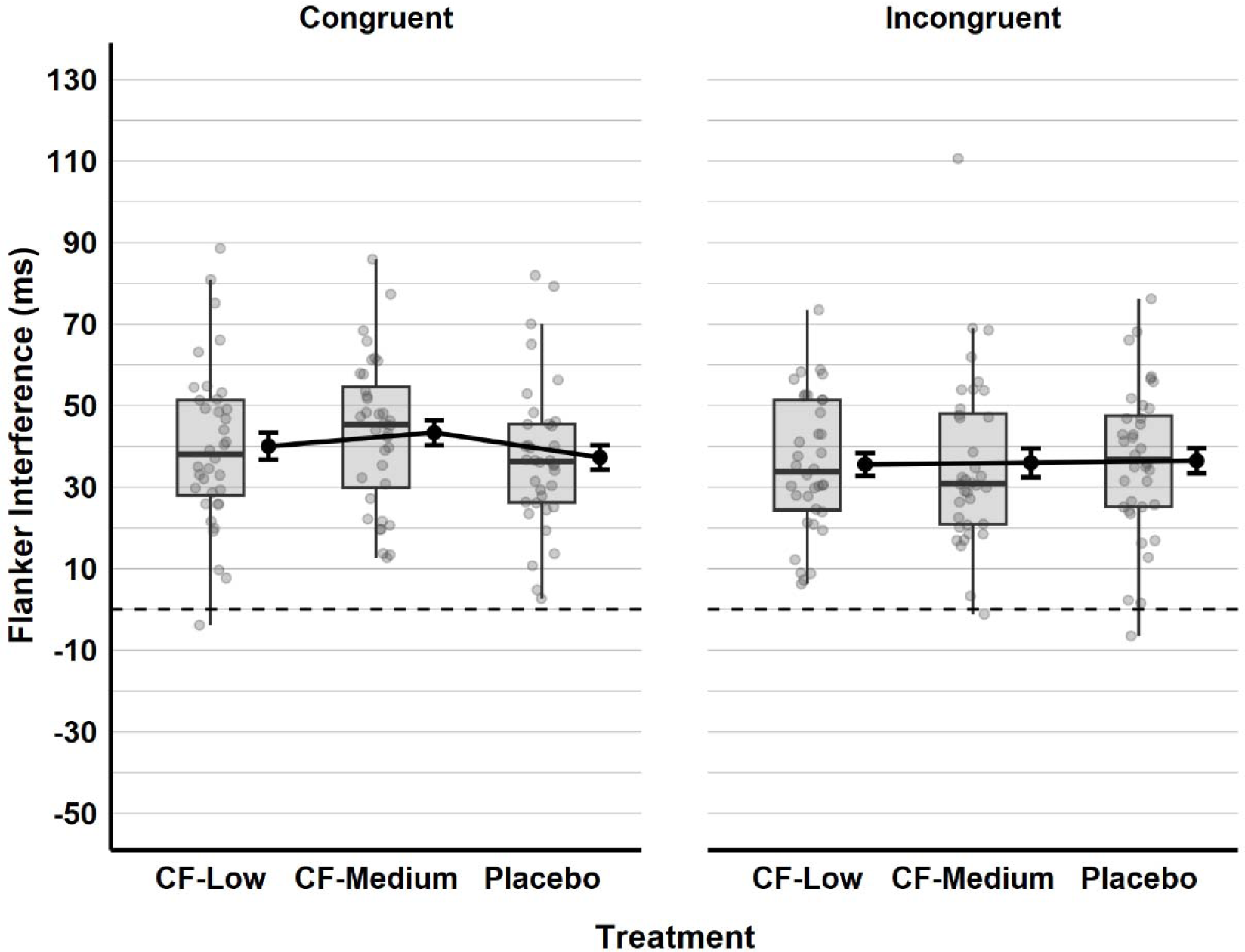
The Flanker Interference Effect on Reaction Time by Previous Trial Congruency and Treatment. *Note*. The boxplots display the data quartiles. Black dots indicate mean values with accompanying error bars representing the standard errors. Grey points mark the individual observations, in each Treatment condition.

#### Simon Task - Accuracy

The acute effects of CF on cognitive control, specifically regarding Simon task accuracy, were evaluated using a Generalised Linear Mixed Models (GLMM) analysis. Initially, random intercepts for subjects was included before adding fixed effects, significantly improving the model. In the null model (without fixed and random effects), the deviance was 15,996 (BIC = 16007), which was reduced to 15,615 (BIC = 15636) upon incorporating random intercepts for subjects. This change in deviance was statistically significant 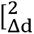 = 380.98, *df* = 1, *p* < .001, *BF_10_* > 100], indicating the necessity of including random intercepts for subjects.

The fixed effects for task conditions (congruent and incongruent) 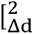 = 94.19, *df* = 2, *p* < .001, *BF_10_* > 100, BIC = 15232] significantly enhanced the model. However, the addition of session 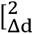 = 4.51, *df* = 2, *p* = .105, *BF_10_* = 0002, BIC = 15249] and Treatment 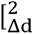 = 2.95, *df* = 2, *p* = .229, *BF_10_* = .0001, BIC = 15251] did not. Adding random slopes for the task conditions 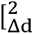 = 31.39, *df* = 2, *p* < .001, *BF_10_* = 151.69, BIC = 15222] also made a significant improvement in the model. The fixed and random effects in the final model are presented in Table 4, and the accuracy by condition and Treatment is given in Figure 6.

**Figure 6.**
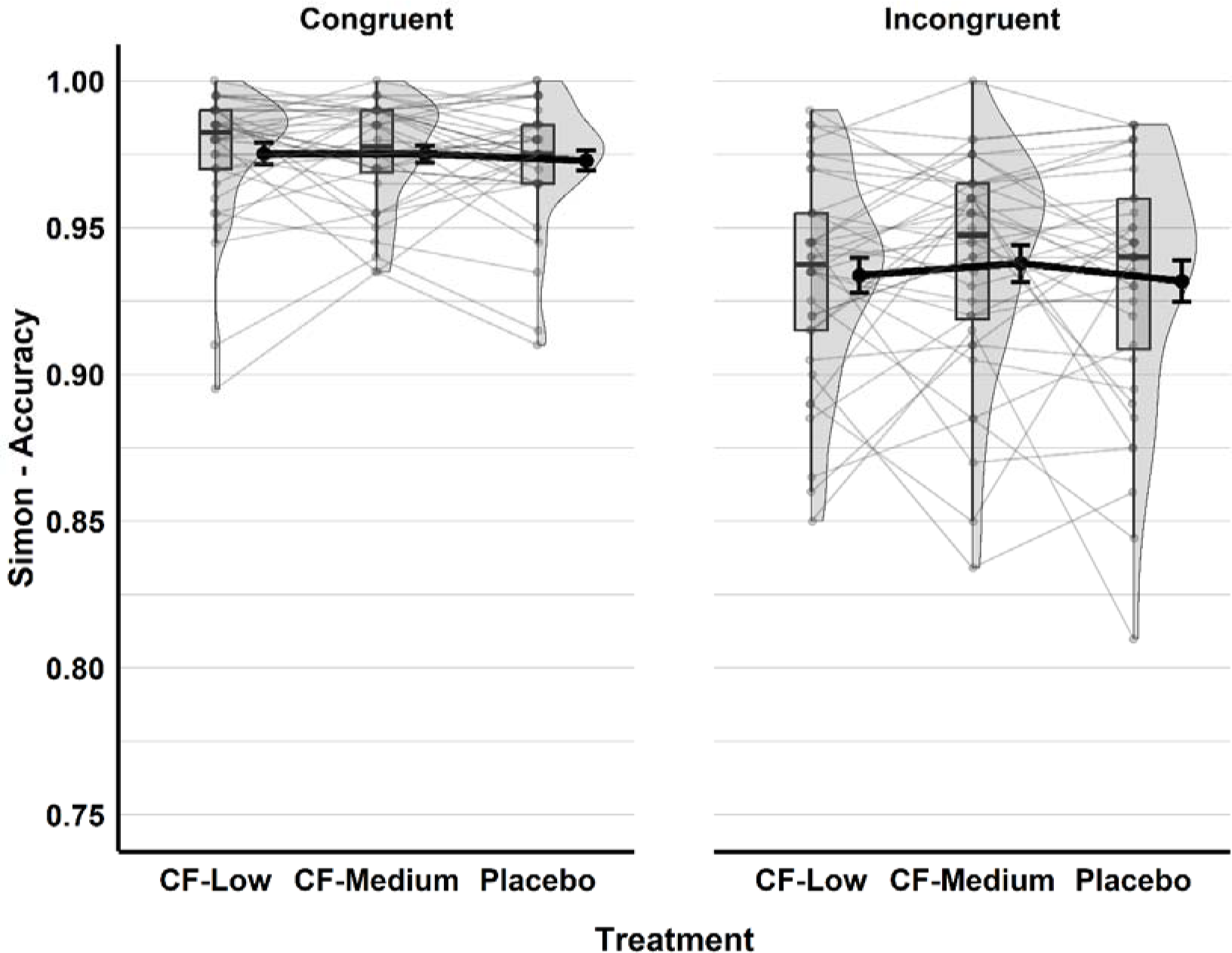
Accuracy in the Simon Task Displayed by Treatment and Condition. *Note*. The boxplots display the data quartiles. Black dots indicate mean values with accompanying error bars representing the standard errors. Grey points mark the individual observations, with lines connecting these points between Treatment conditions.

**Table 4.**
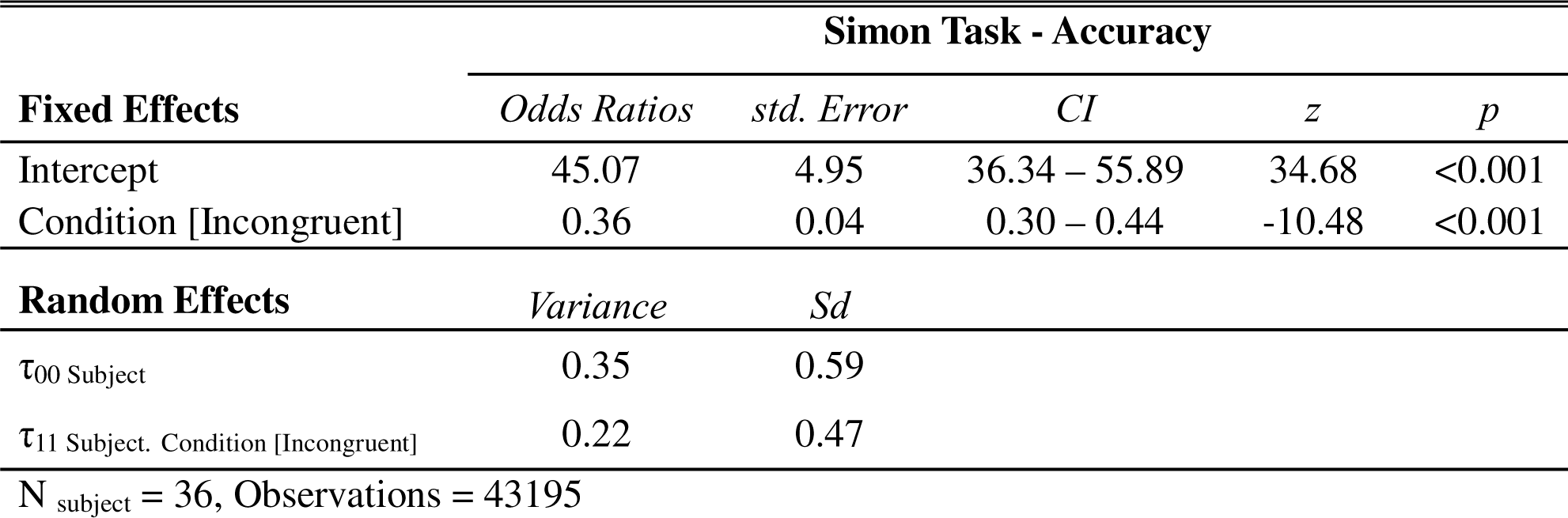
Generalised Linear Mixed Model Results for the Effects of CF on Accuracy in the Simon Task.

| Simon Task - Accuracy |  |  |  |  |  |
| --- | --- | --- | --- | --- | --- |
| Fixed Effects | *Odds Ratios* | *std. Error* | *CI* | *z* | *p* |
| Intercept | 45.07 | 4.95 | 36.34 – 55.89 | 34.68 | <0.001 |
| Condition [Incongruent] | 0.36 | 0.04 | 0.30 – 0.44 | -10.48 | <0.001 |
| Random Effects | *Variance* | *Sd* |  |  |  |
| τ<sub>00</sub> Subject | 0.35 | 0.59 |  |  |  |
| τ<sub>11</sub> Subject. Condition [Incongruent] | 0.22 | 0.47 |  |  |  |
| N<sub>subject</sub> = 36, Observations = 43195 |  |  |  |  |  |

Tukey’s pairwise comparisons revealed that accuracy was significantly higher in the congruent condition (*prob.* = .978, SE = .002, 95% CL [.973 – 0.984]), compared to the incongruent condition (*prob.* = .942, SE = .005, 95% CL [.930 – 0.952]), with *Z* = 10.45, *p* < .001. Nevertheless, the inclusion of participants’ gender 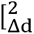 = .18, *df* = 1, *p* = .672, *BF_10_* = .005, BIC = 15233] and BMI scores 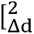 = .001, *df* = 1, *p* = .975, *BF_10_* = .005, BIC = 15233] as predictors did not significantly improve the model.

#### Simon Task – Reaction Time Analysis (General)

In the Linear Mixed Model (LMM) analysis concerning response times for correct trials in the Simon task, the initial deviance without any fixed or random effects was 483852 (BIC = 483874). The introduction of random intercepts for participants decreased the deviance to 479120 (BIC = 479152), which was statistically significant 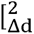 = 4732.8, *df* = 1, *p* < .001, *BF_10_* > 100]. Fixed effects for the task conditions 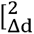 = 1634.8, *df* = 1, *p* < .001, *BF_10_* > 100, BIC = 477527] and session 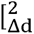 = 637.95, *df* = 2, *p* < .001, *BF_10_* > 100, BIC= 476911] significantly improved the model, while adding the Treatment 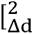 = 9.995, *df* = 2, *p* = .030, *BF_10_*= .004, BIC = 476925] did not yield a significant improvement. After the inclusion of fixed effects, the model was significantly enhanced by adding random slopes for the task conditions 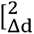 = 199.2, *df* = 2, *p* < .001, *BF_10_* > 100, BIC = 476723] and sessions 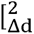 = 437.6, *df* = 7, *p* < .001, *BF_10_* > 100, BIC = 476360]. Table 5 displays the final model results, and Figure 7 illustrates the response times for the Simon task according to the different conditions and Treatments.

**Figure 7.**
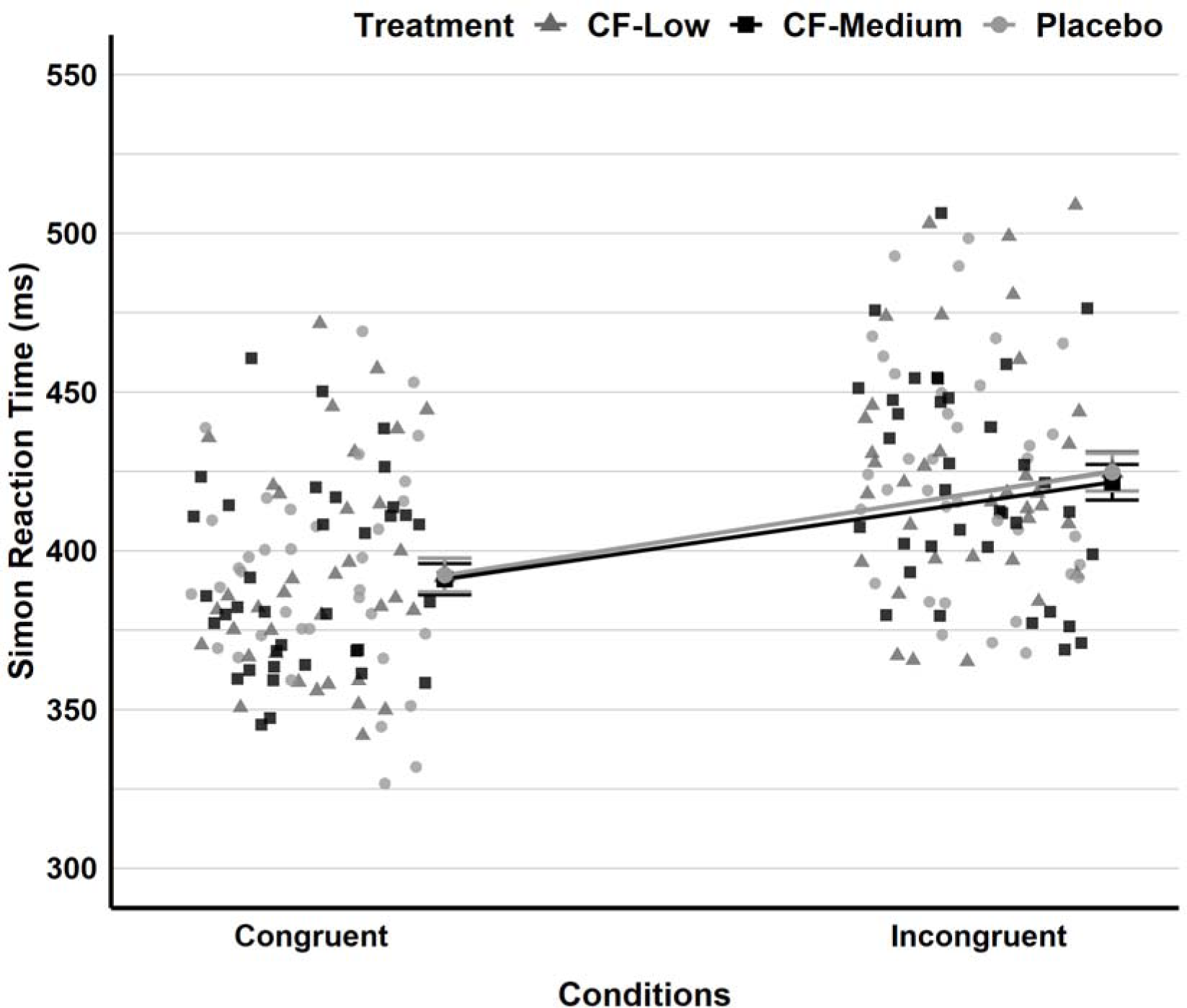
Response Times in the Simon Task across Treatment and Conditions. *Note*. The Placebo condition is denoted by light grey circles for individual observations and larger dots for means and standard errors. Similarly, medium-light grey triangles indicate the CF-Low condition, while the CF-Medium condition is depicted with black squares. Lines connect the group means.

**Table 5.**
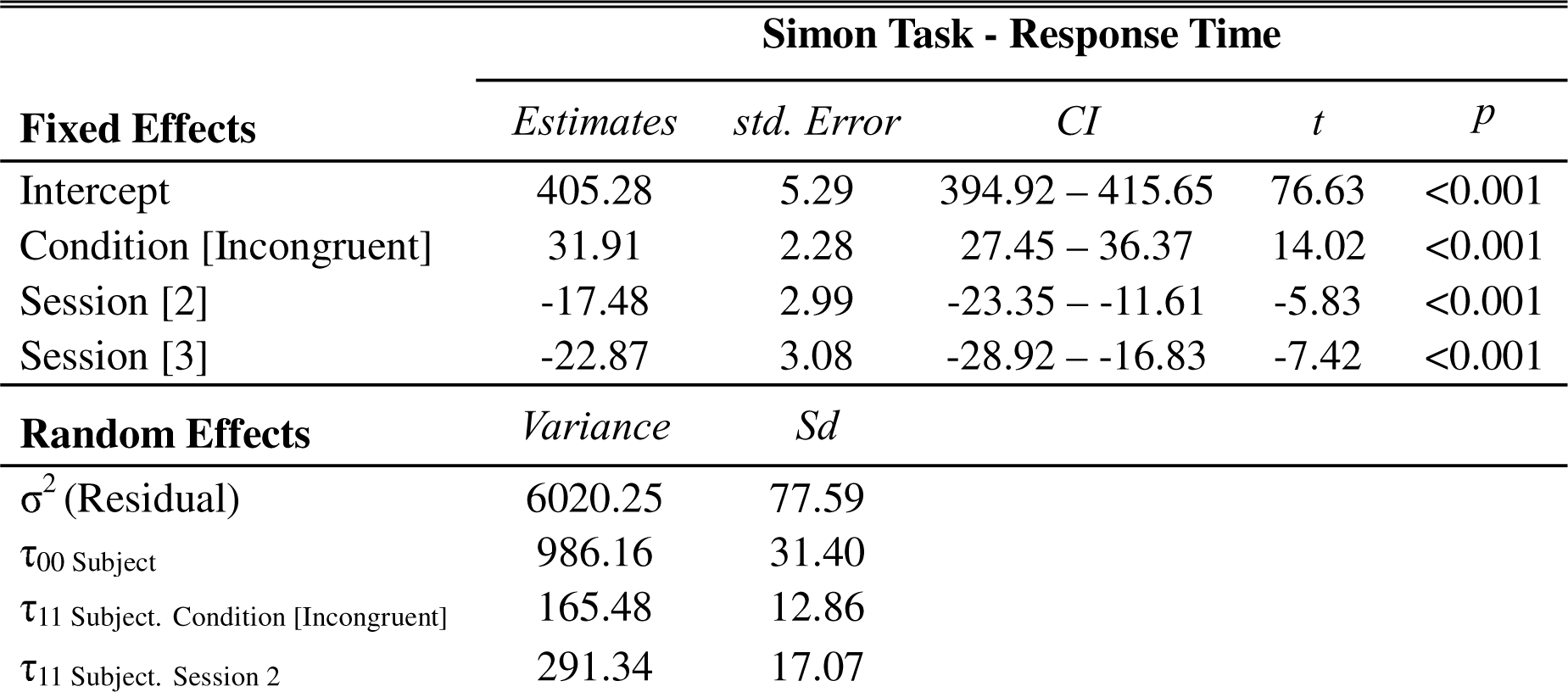
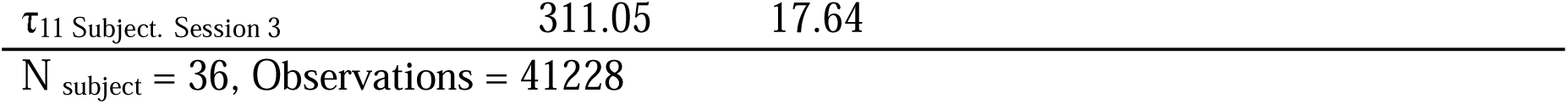
Linear Mixed Model Results for the Effects of CF on Response Times in the Simon Task.

| **Simon Task - Response Time** |  |  |  |  |  |
| --- | --- | --- | --- | --- | --- |
| **Fixed Effects** | *Estimates* | *std. Error* | *CI* | *t* | *p* |
| Intercept | 405.28 | 5.29 | 394.92 – 415.65 | 76.63 | <0.001 |
| Condition [Incongruent] | 31.91 | 2.28 | 27.45 – 36.37 | 14.02 | <0.001 |
| Session [2] | -17.48 | 2.99 | -23.35 – -11.61 | -5.83 | <0.001 |
| Session [3] | -22.87 | 3.08 | -28.92 – -16.83 | -7.42 | <0.001 |
| **Random Effects** | *Variance* | *Sd* |  |  |  |
| $\sigma^2$ (Residual) | 6020.25 | 77.59 | | | |
| $\tau_{00}$ Subject | 986.16 | 31.40 | | | |
| $\tau_{11}$ Subject. Condition [Incongruent] | 165.48 | 12.86 | | | |
| $\tau_{11}$ Subject. Session 2 | 291.34 | 17.07 | | | |

Tukey pairwise comparisons indicated that response times were significantly faster for congruent trials (*Estimated marginal mean* = 392, SE = 4.71, 95% CI [383 – 401]) than for incongruent trials (*Estimated marginal mean* = 424, SE = 5.30, 95% CI [413 – 434]), with Z = −14.02, *p* < .001. There was a gradual decrease in response times when considering the number of sessions as expected. In the exploratory analysis, however, the inclusion of gender 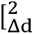 = .71, *df* = 1, *p* = .403, *BF_10_* = .159, BIC = 476386] and BMI scores 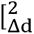 = .63, *df* = 1, *p* = .426, *BF_10_* = .026, BIC = 476386] did not significantly improve the model.

#### Simon Task – Reaction Time Analysis (Sequential)

In addition to the general reaction time analysis in the Simon task, sequential effects based on the trial preceding the current trial were also calculated. Results from a two-way ANOVA indicated that the main effect of trial flow conditions (sequences based on previous trials, incongruent-congruent, iC; congruent-congruent, cC; incongruent-incongruent, iI; and congruent-incongruent, cI) was statistically significant, *F*(3, 420) = 54.78, *p* < .001, 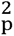 = .28. However, neither the main effects of Treatment, *F*(2, 420) = 0.21, *p* = .813, 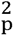 = .001, nor the interaction of Treatment and trial conditions, *F*(6, 420) = 0.14, *p* = .991, 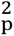 = .002, reached statistical significance (see Figure 8).

**Figure 8.**
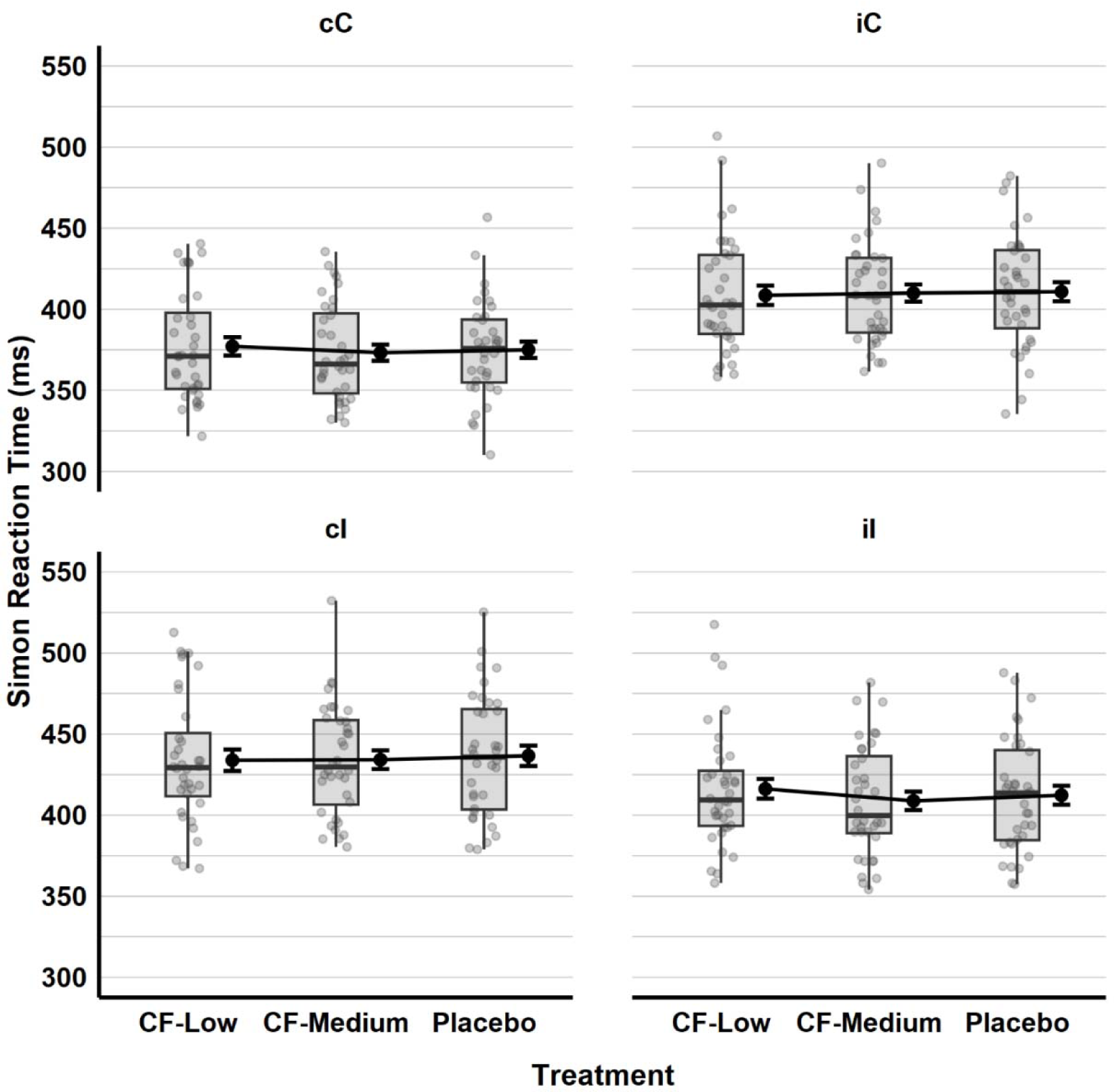
Reaction times in the Simon Task Across Previous Trial Congruency and Treatment. *Note*. Reaction times in previous (lower case) and current (upper case) trial conditions. The boxplots display the data quartiles. Black dots and lines indicate mean values with accompanying error bars representing the standard errors. Grey points mark the individual observations in each Treatment condition.

In pairwise comparisons, post-hoc analysis using Tukey’s HSD indicated a significant difference in reaction times for Congruent current trials based on the type of preceding trial. Specifically, when the current Congruent trial followed another Congruent trial (cC), reaction times were significantly faster compared to when the current Congruent trial followed an Incongruent trial (iC), 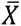Δ = 34.72, 95% CI [22.55, 46.89], *p* < .001. Similarly, for Incongruent current trials, reaction times were significantly faster when following a Incongruent trial (iI), compared to those following an Congruent trial (cI), 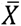Δ = 22.45, 95% CI [34.62, 10.29], *p* < .001. These findings highlight the impact of trial sequence on the reaction times, independent of the effects of CF.

#### Simon Effect on Reaction Time (General)

We calculated the Simon effect on reaction time, indicating the difference between the mean reaction times for incongruent and congruent trials for each subject. Two-way ANOVA analysis revealed no significant main effects for Treatment, *F*(2, 99) = 0.22, *p* = .80, 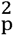 = .004, and Session, *F*(2, 99) = 1.47, *p* = .24, 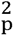 = .03. The interaction of Treatment and Session was statistically significant, *F*(4, 99) = 2.91, *p* = .03, 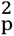 = .11. Post-hoc tests showed that this only concerned a decrease of the Simon effect within the low CF condition from the first to the third session, 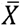 = −23.32, 95% *CI* [−44.14, −2.50], *p* = .017.

#### Simon Effect on Reaction Time (Sequential)

Similar to the sequential effects of previous trials on general reaction time, we also examined the impact of the previous trial on Simon interference effects on reaction time. A two-way ANOVA revealed that the main effects of previous conditions (congruent or incongruent) were statistically significant, *F*(1, 210) = 357.93, *p* < .001, 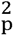 = .63. However, neither the main effects of Treatment, *F*(2, 210) = 0.21, *p* = .815, 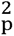 = .002, nor the interaction of Treatment and conditions, *F*(2, 210) = 1.80, *p* = .167, 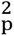 = .002, were significant (see Figure 9).

**Figure 9.**
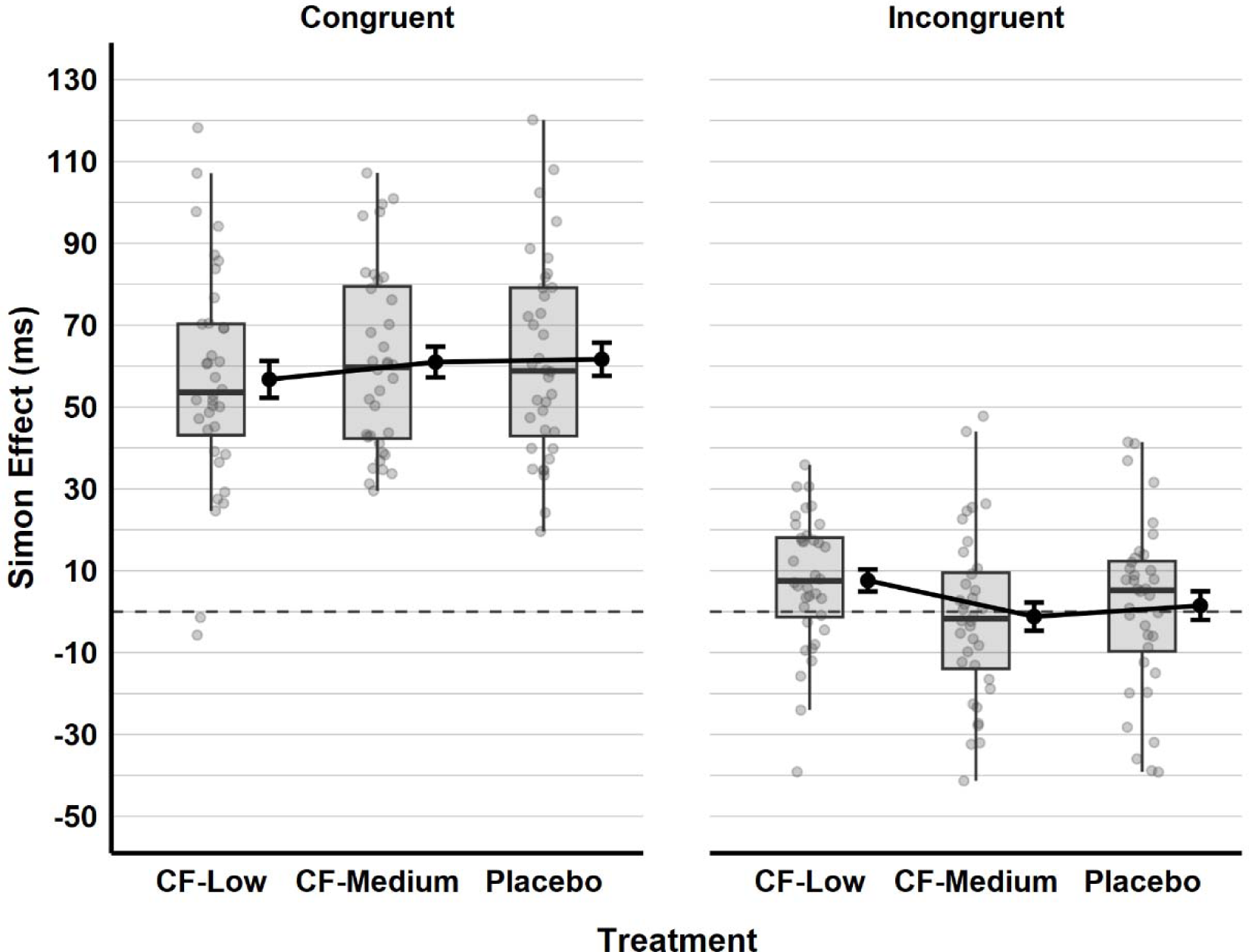
The Simon Effect on Reaction Times by Treatment and Type of Previous Trial. *Note*. The boxplots display the data quartiles. Black dots indicate mean values with accompanying error bars representing the standard errors. Grey points mark the individual observations in each Treatment condition.

Post hoc comparisons using Tukey’s HSD test indicated that the Simon effect on reaction times for trials following incongruent stimulus was significantly lower compared to when the previous trial was congruent, 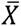 Δ = −57.18, 95% *CI* [−63.13, −51.22], *p* < .001.

### Response Inhibition

#### Go/No-go Task – Commission Error

In evaluating the performance in the Go/No-go task, both commission and omission errors were assessed, as well as reaction time for correct go trials. The acute effects of CF on commission errors were assessed through a GLMM analysis. The model initially included random intercepts for subjects before fixed effects were added, which significantly improved the model with deviance changing from 12,436 (BIC = 12445) to 12,027 (BIC = 12046), 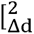 = 408.68, *df* = 1, *p* < .001, *BF_10_* > 100], suggesting the need to include random intercepts for subjects. In the analysis of fixed effects, neither session 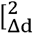 = .58, *df* = 2, *p* = .747, *BF_10_* = .0009, BIC = 12065] nor Treatment 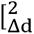 = 4.16, *df* = 2, *p* = .125, *BF_10_* = .0005, BIC = 12061] improved the model significantly. Consequently, the random slope analysis stage was omitted due to the absence of significant fixed effects in the previous stage. Also, the gender of participants 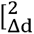 = 5.21, *df* = 1, *p* = .022, *BF_10_* = .113, BIC = 12050] and BMI scores 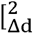 = 1.41, *df* = 1, *p* = .234, *BF_10_* = .017, BIC = 12054] did not significantly improve the model. Commission error percentages by Treatment are given in Figure 10.

**Figure 10.**
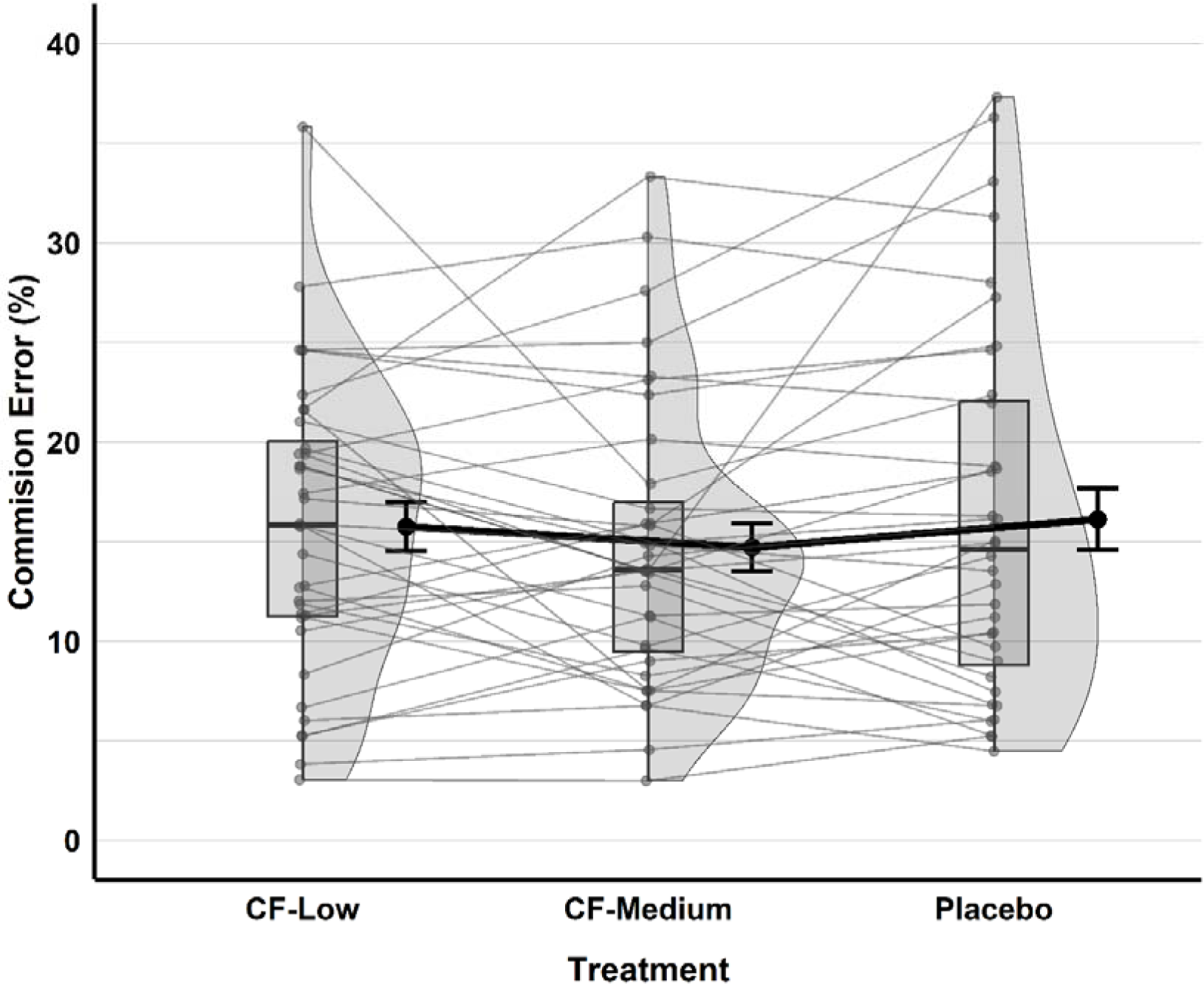
Percentage of Commission Errors in the Go/No-go Task across Treatment. *Note*. The boxplots display the data quartiles. Mean values are indicated by black dots with accompanying error bars representing the standard errors. Grey points mark the individual observations, with lines connecting these points between Treatment conditions.

#### Go/No-go Task – Omission Error

The acute effects of CF on omission errors were evaluated with a GLMM analysis. At the start, random intercepts for each subject were included in the model, and the model improved significantly 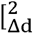 = 408.68, *df* = 1, *p* < .001, *BF_10_* > 100]. In the null model, the deviance was 2573 (BIC = 2583), which was reduced to 2408 (BIC = 2429) by adding random intercepts for subjects, indicating the necessity of including random intercepts for subjects.

Similar to commission errors, there was no evidence that session 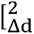 = 9.53, *df* = 2, *p* = .009, *BF_10_* = .004, BIC = 2440] and Treatment 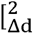 = 7.98, *df* = 2, *p* = .019, *BF_10_* = .002, BIC = 2441] had a significant effect on improving the model. Likewise, gender 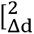 = 4.22, *df* = 1, *p* = .039, *BF_10_* = .049, BIC = 2435] and BMI scores 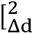 = .11, *df* = 1, *p* = .745, *BF_10_* = .006, BIC = 2439] did not significantly enhance the model either. Omission error percentages by Treatment are presented in Figure 11.

**Figure 11.**
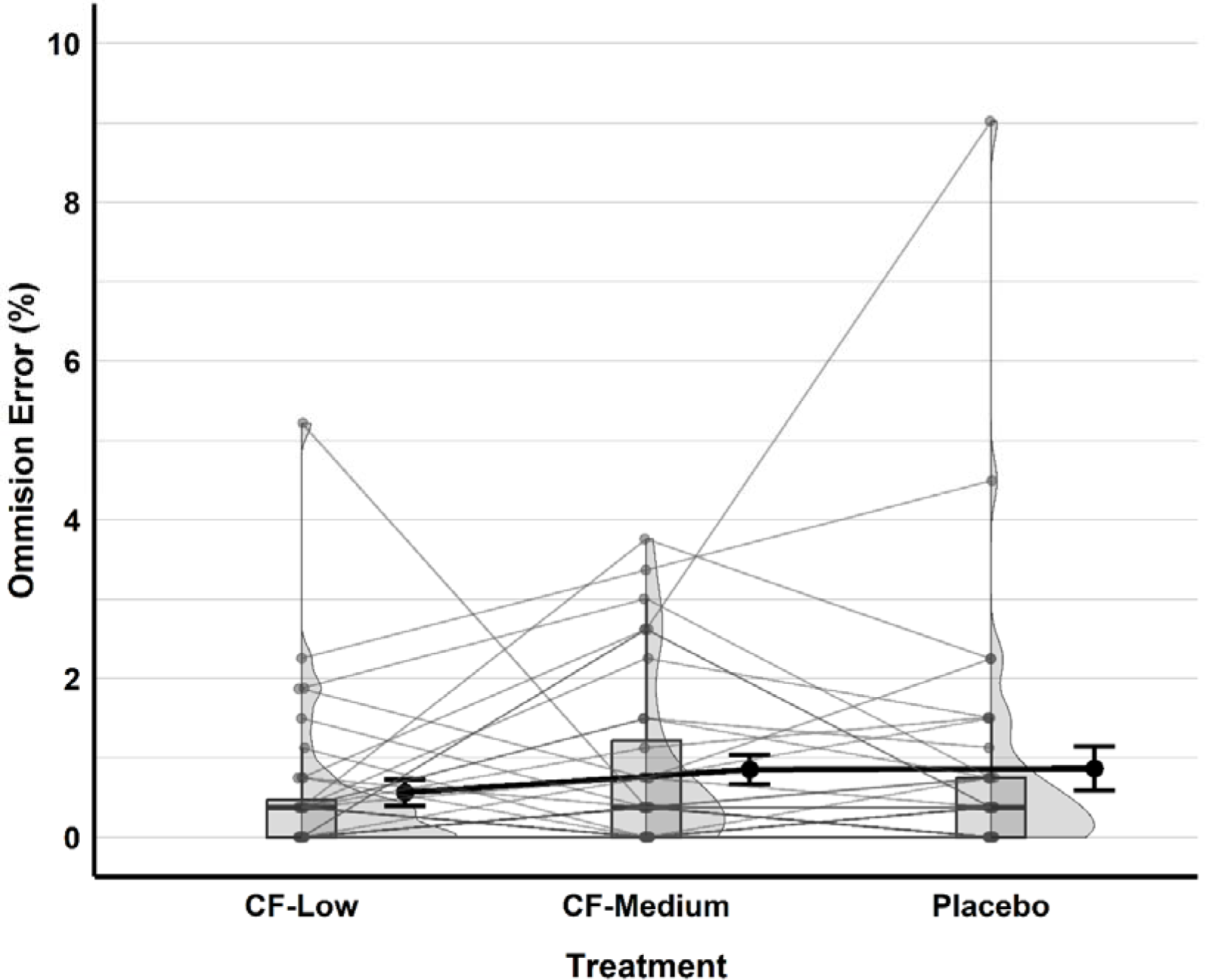
Percentage of Omission Errors in the Go/No-go Task across Treatment. *Note*. The boxplots display the data quartiles. Mean values are indicated by black dots with accompanying error bars representing the standard errors. Grey points mark the individual observations, with lines connecting these points between Treatment conditions.

#### Go/No-go Task – Reaction Time

In assessing the acute effects of CF on response times for Go trials in the Go/No-go task, Linear Mixed Model analysis was employed. Firstly, the model’s deviance was 344472 (BIC = 344472), and it decreased to 339798 (BIC = 339829) after the inclusion of random intercepts for subjects 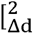 = 4673.7, *df* = 1, *p* < .001, *BF_10_* > 100].

In the analysis of fixed effects, the session 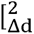 = 82.5, *df* = 2, *p* < .001, *BF_10_* > 100, BIC = 339767], Treatment 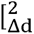 = 31.7, *df* = 2, *p* < .001, *BF_10_* = 2520.4, BIC = 339756] and the interaction of session and Treatment 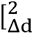 = 134.0, *df* = 4, *p* < .001, *BF_10_* > 100, BIC = 339663] improved the model significantly.

After fixed effects, the model was further improved by introducing random slopes for the session 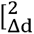 = 1134.1, *df* = 5, *p* < .001, *BF_10_* > 100, BIC = 338547], while the addition of Treatment as a random slope 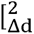 = 13.2, *df* = 9, *p* = .152, *BF_10_*< .0001, BIC = 338626] did not yield a significant enhancement. The final model results are reported in Table 6.

**Table 6.**
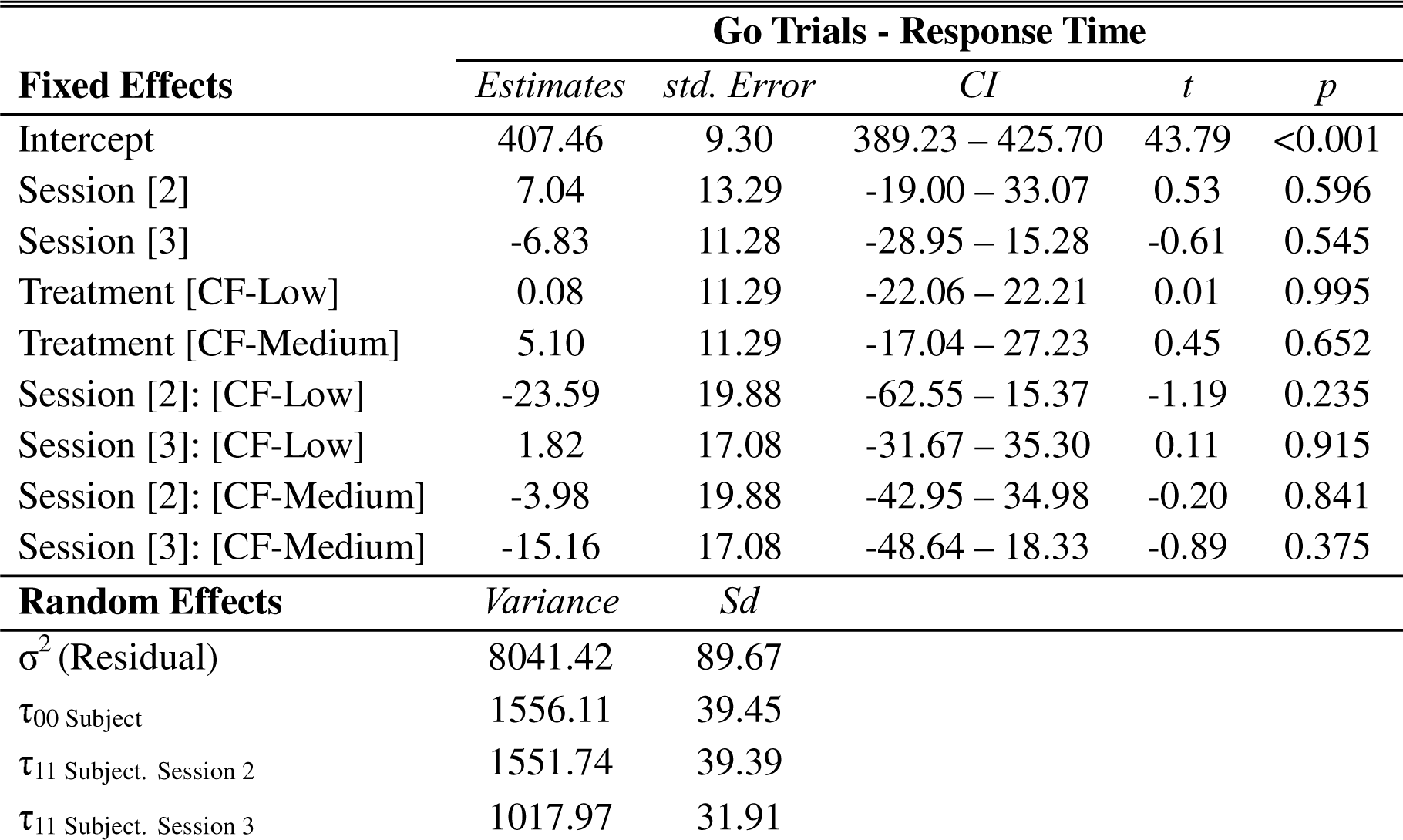
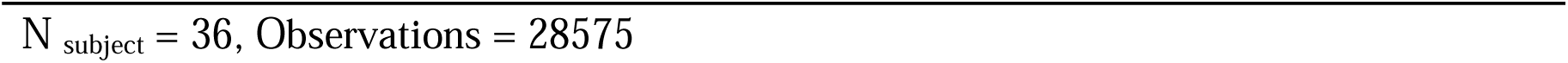
Linear Mixed Model Results for the Effects of CF on Response Time in the Go/No-go Task.

| **Fixed Effects** | **Go Trials - Response Time** |  |  |  |  |
| --- | --- | --- | --- | --- | --- |
|  | *Estimates* | *std. Error* | *CI* | *t* | *p* |
| Intercept | 407.46 | 9.30 | 389.23 – 425.70 | 43.79 | <0.001 |
| Session [2] | 7.04 | 13.29 | -19.00 – 33.07 | 0.53 | 0.596 |
| Session [3] | -6.83 | 11.28 | -28.95 – 15.28 | -0.61 | 0.545 |
| Treatment [CF-Low] | 0.08 | 11.29 | -22.06 – 22.21 | 0.01 | 0.995 |
| Treatment [CF-Medium] | 5.10 | 11.29 | -17.04 – 27.23 | 0.45 | 0.652 |
| Session [2]: [CF-Low] | -23.59 | 19.88 | -62.55 – 15.37 | -1.19 | 0.235 |
| Session [3]: [CF-Low] | 1.82 | 17.08 | -31.67 – 35.30 | 0.11 | 0.915 |
| Session [2]: [CF-Medium] | -3.98 | 19.88 | -42.95 – 34.98 | -0.20 | 0.841 |
| Session [3]: [CF-Medium] | -15.16 | 17.08 | -48.64 – 18.33 | -0.89 | 0.375 |
| **Random Effects** | *Variance* | *Sd* |  |  |  |
| $\sigma^2$ (Residual) | 8041.42 | 89.67 | | | |
| $\tau_{00}$ Subject | 1556.11 | 39.45 | | | |
| $\tau_{11}$ Subject. Session 2 | 1551.74 | 39.39 | | | |
| $\tau_{11}$ Subject. Session 3 | 1017.97 | 31.91 | | | |

For the final model, with random slopes of session, Tukey pairwise comparisons indicated that response times for correct Go trials in the Go/No-go task were changed neither for sessions nor Treatment conditions. It should be noted that the fixed effects of Treatment and its interaction with Session become non-significant after the inclusion of random slopes for Session. This suggests there is considerable variability in the effects of Treatment on reaction times across different sessions. Also, the exploratory analysis showed that the inclusion of gender 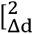 = .58, *df* = 1, *p* = .447, *BF_10_* = .252, BIC = 338439] and BMI scores 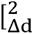 = .003, *df* = 1, *p* = .957, *BF_10_* = .031, BIC = 338613] did not significantly improve the model. Figure 12 illustrates the response times for the Go trials in the Go/No-go task according to different Treatments and sessions.

**Figure 12.**
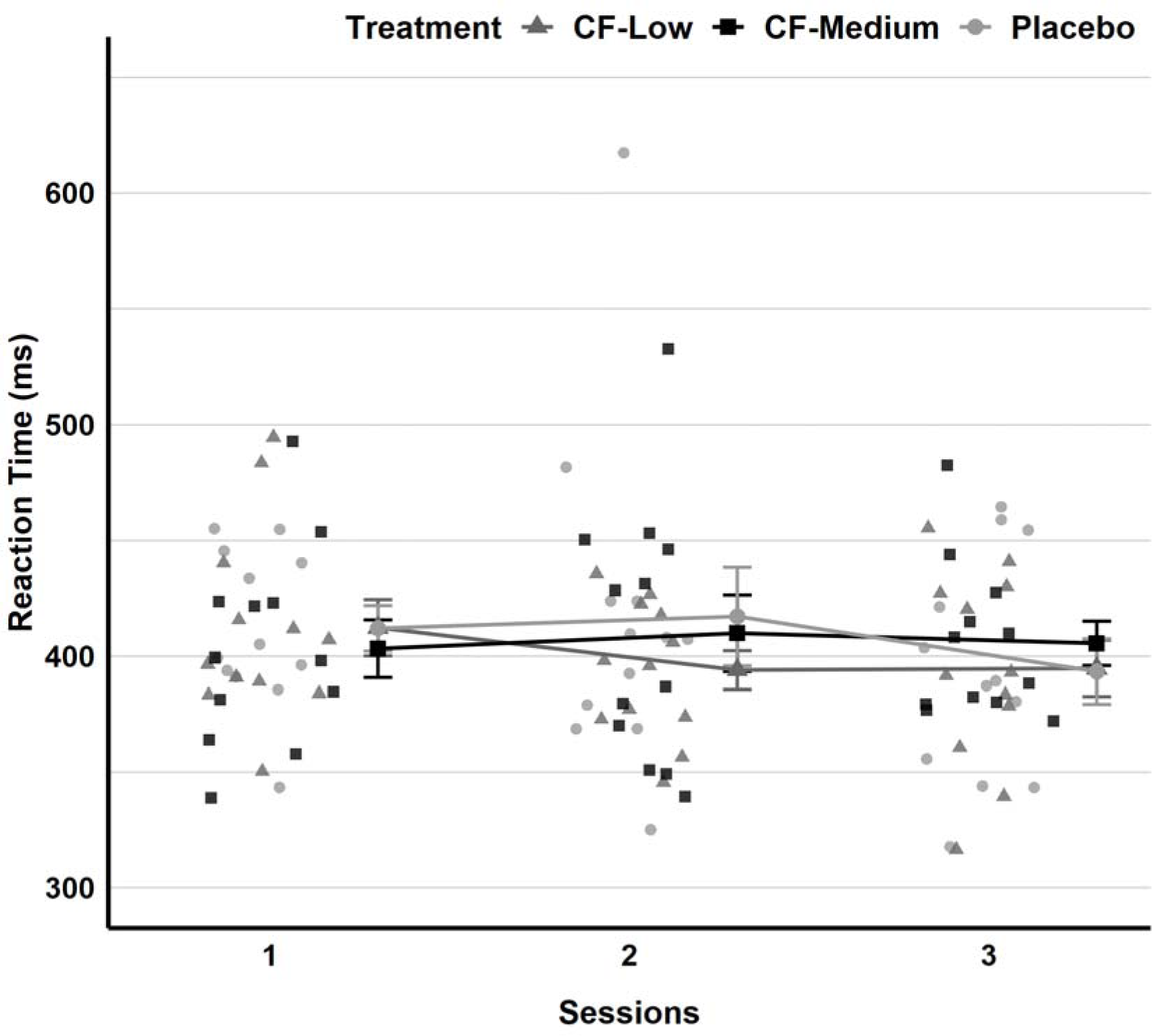
Response Times in Go/No-go Task across Sessions and Treatment. Note. The Placebo condition is denoted by light grey circles for individual observations and larger dots for means and standard errors. The CF-Low condition is indicated by medium-light grey triangles, while the CF-Medium condition is depicted with black squares.

## Discussion

In this pre-registered, randomized, placebo-controlled, gender-balanced, double-blind, and crossover-designed study, we investigated the acute effects of cocoa flavanols (CF), at doses of 415mg and 623mg, on cognitive control and response inhibition in healthy young adults. While previous research has examined the acute effects of CF on cognitive control, our study uniquely applied the Simon and Go/No-go tasks, along with the Flanker task, to provide a comprehensive evaluation of CF’s impact on cognitive control and response inhibition. Our analysis revealed that neither the 415mg nor the 623mg doses of CF significantly affected cognitive control and response inhibition. Specifically, in both the Flanker and Simon tasks, CF intake did not influence accuracy, general reaction time, or interference levels. Furthermore, a sequential analysis, accounting for the effects of previous trials in the Simon and Flanker tasks, showed no detectable improvements attributable to CF intake. Additionally, the Go/No-go task measures indicated no significant changes in commission and omission errors, nor in reaction times.

These results, particularly for the Flanker task, diverge from those reported previously. Specifically, Decroix and colleagues (2019) observed that an acute intake of 900mg CF improved overall reaction time, regardless of the interaction between treatment and task conditions. The same study also found that this dosage of CF increased BOLD responses in the supramarginal gyrus of the parietal lobe and the inferior frontal gyrus in both diabetic patients and healthy controls. However, aligning with our study’s findings, they did not observe any improvement in accuracy and Flanker interference. Unlike our study, Decroix et al. conducted their research with a high dose of CF, and relatively small sample sizes, including 11 participants each in the Type-1 diabetes and matched healthy control groups, all of whom were middle-aged. The discrepancies between our results and theirs could thus stem from these variations in dosage, age, and the clinical characteristics of the participants.

Regarding the effects of different doses of CF on cognitive functions, previous studies (Boolani et al., 2017; Grassi et al., 2016; Karabay et al., 2018; Massee et al., 2015; Scholey et al., 2010) have reported cognitive performance improvements mostly with small to medium doses (250-720mg), rather than high doses (747-994mg). Additionally, a meta-analysis by Barrera-Reyes et al. (2020) concluded that there is a medium to large effect size for the effects of CF on memory and executive function with intermediate doses (500-750mg) of CF. In our study, we tested the effects of 415mg and 623mg CF, corresponding to small to medium doses, and observed no improvements in healthy young adults. Conversely, the significant improvements in Flanker task reaction times with a 900mg CF dose in a middle-aged sample that were found previously by Decroix and colleagues (2019) suggest that higher doses may be more effective in older adults.

Moreover, considering the characteristics of the sample studied by Decroix and colleagues (2019), Type-1 diabetes, described as a chronic autoimmune disease characterized by insulin deficiency and resistance (Cade, 2008), can result in cognitive and neurovascular dysfunctions (Brands et al., 2005; Lasta et al., 2013). Although no significant interaction effects between group (diabetes and healthy control) and treatment conditions were found, the interaction of group and task condition was significant, with healthy controls showing slower RTs for incongruent versus congruent and neutral tasks. However, individuals with diabetes showed no significant differences in RT across task types and exhibited a smaller flanker interference effect compared to healthy controls. This suggests that cognitive processing might differ between healthy individuals and those with diabetes, with the healthy group showing more sensitivity to task condition changes. Taken together, the effects of CF in healthy young adults may differ from those in middle-aged groups with these specific characteristics.

Consistent with the findings of our study, longitudinal research (Brickman et al., 2023; L. K. Yeung et al., 2023) demonstrated that a three-year intervention with 500mg CF did not significantly impact Flanker task performance in older adults. However, in contrast to this result, studies on children using an anthocyanin (253mg) from wild blueberries—a different subclass of flavonoids—showed an acute increase in accuracy during incongruent trials in the Flanker task, though not in reaction times (Barfoot et al., 2019; Whyte et al., 2016, 2017). These outcomes suggest that while the acute and long-term effects of CF on cognitive control appear limited in older and young adults.

Aligned with our findings on the Flanker task, this study observed no significant enhancement in Simon task performance following two different low-to-medium doses of CF. The Simon task, as noted by Mansfield et al. (2013), engages more visual-spatial cognitive control processes compared to the Flanker task. Despite the absence of direct evidence on CF’s effects on Simon task performance, prior research has demonstrated benefits of acute CF administration on spatial attention efficiency, evidenced by faster reaction times (Karabay et al., 2018), and improved visual spatial working memory accuracy in location tasks (Field et al., 2011) among young adults. Furthermore, an fMRI study (Francis et al., 2006) showed that CF increased activation in the right posterior parietal cortex, crucial for spatial perception and visuomotor control (Malhotra et al., 2009; Rushworth et al., 2001). Similarly, a functional NIRS study (Decroix et al., 2018) observed acute CF-induced activation increases in the right prefrontal cortex, associated with higher cognitive functions such as decision-making and problem-solving (Friedman & Robbins, 2022; Kolk & Rakic, 2022). However, not all of these studies reported commensurate behavioural improvements in task-switching (Francis et al., 2006) or basic spatial object recognition (Decroix et al., 2018). It is conceivable that the influence of CF may be limited to more complex processes beyond basic cognitive control. Further research may attempt to compare this more systematically.

In contrast to our study, in which we did not observe improvements in reaction times or in omission and commission errors, Field and colleagues (2011) noted reduced reaction times during predictable trials of a task that arguably also required response inhibition with 720mg CF. This task required participants to identify presented letters (’X’ or ‘Y’) or digits by pressing labelled keys—’X’ and ‘Y’ for letters on the screen and ‘N’ for digits. Enhancements were observed only in the predictable phase, which required comparatively less inhibitory effort for letter identification. Conversely, the second phase, which introduced digits and demanded higher inhibitory effort, showed no significant effect. Additionally, the gender imbalance in Field et al.’s study (22 female, 8 male), in contrast to our gender-balanced research, suggests that there could be gender differences in the effects of CF, also in view of the absence of interactions with BMI in our study. Similar to our results, Lamport et al. (2016) and Kean et al. (2015) found no acute or long-term improvements in Go/No-go task performance with flavonoid-rich interventions in young and older adults, respectively.

At this point, it may be noted that the current study is subject to several limitations. Notably, it lacks physical/cardiovascular evaluations, such as saliva sampling or heart rate (HR) variability measurements post-CF intake, to confirm the presence of CF in metabolism. Additionally, the effects of CF were assessed on cognitive tasks at a single point in time; a longitudinal approach could provide a more comprehensive understanding of its effects. The age range of our sample, consisting of young adults from various fields with presumably higher cognitive abilities, may have made it challenging to detect significant manipulations, although task performance was not consistently at ceiling. Investigating CF’s impact across different age groups could elucidate its developmental role and potentially uncover effects in older adults. Furthermore, the study utilized basic versions of cognitive tasks; a comparison with more complex task versions could reveal how CF influences higher-level cognitive control, under varying levels of demand.

Despite these limitations, the study’s pre-registration, as well as its randomized, double-blind, gender-balanced, and crossover design strengthens its capacity to detect potential effects while controlling for individual differences. Moreover, we employed a multilevel model approach, accounting for individual variations in task performance with random intercepts and random slopes. Ultimately, this contributes to the strength of the present evidence that 415mg and 623mg doses of CF do not acutely enhance cognitive functions as measured by the Flanker, Simon, and Go/No-go tasks.

## Notes

### Competing Interest Statement

The authors have declared no competing interest.

